# Deciphering the quality of SARS-CoV-2 specific T-cell response associated with disease severity, immune memory and heterologous response

**DOI:** 10.1101/2021.12.28.474325

**Authors:** Alberto Pérez-Gómez, M Carmen Gasca-Capote, Joana Vitallé, Francisco J. Ostos, Ana Serna-Gallego, María Trujillo-Rodríguez, Esperanza Muñoz-Muela, Teresa Giráldez-Pérez, Julia Praena-Segovia, María D. Navarro-Amuedo, María Paniagua-García, Manuel García-Gutiérrez, Manuela Aguilar-Guisado, Inmaculada Rivas-Jeremias, María R. Jimenez-Leon, Sara Bachiller, Alberto Fernández-Villar, Alexandre Pérez-González, Alicia Gutiérrez-Valencia, Mohammed Rafii-El-Idrissi Benhnia, Daniela Weiskopf, Alessandro Sette, Luis F. López-Cortes, Eva Poveda, Ezequiel Ruiz-Mateos, Virgen del Rocío Hospital COVID-19, COHVID-GS Working Teams

## Abstract

SARS-CoV-2 specific T-cell response has been associated with disease severity, immune memory and heterologous response to endemic coronaviruses. However, an integrative approach combining a comprehensive analysis of the quality of SARS-CoV-2 specific T-cell response with antibody levels in these three scenarios is needed. In the present study we found that, in acute infection, while mild disease was associated with high T-cell polyfunctionality biased to IL-2 production and inversely correlated with anti-S IgG levels, combinations only including IFN-γ with absence of perforin production predominated in severe disease. Seven months after infection, both non-hospitalized and previously hospitalized patients presented robust anti-S IgG levels and SARS-CoV-2 specific T-cell response. In addition, only previously hospitalized patients showed a T-cell exhaustion profile. Finally, combinations including IL-2 in response to S protein of endemic coronaviruses, were the ones associated with SARS-CoV-2 S-specific T-cell response in pre-COVID-19 healthy donors’ samples. These results have implications for protective immunity against SARS-CoV-2 and recurrent COVID-19 and may help for the design of new prototypes and boosting vaccine strategies.

## INTRODUCTION

Host immune response against Severe Acute Respiratory Syndrome Coronavirus 2 (SARS-CoV-2) infection is a key factor in the progression of Coronavirus Disease 2019 (COVID-19) (1) and its deregulation results in fatal disease in hospitalized COVID-19 patients (2, 3). The coordination of different branches of adaptive immunity, such as CD4+, CD8+ T-cell and antibody responses, is essential for the resolution of COVID-19 (4). Despite the already known role of T-cell response against SARS-CoV-2 infection, there are still gaps that need to be clarify in relation to the quality of this response and its association with: i) disease severity in acute infection, ii) long-lasting immune memory and iii) the heterologous response found in healthy donors (5).

Seminal studies in SARS-CoV-1 infection models showed that both CD4+ (6) and CD8+ (7) T-cell response were involved in protection and virus clearance in acute infection. In SARS-CoV-2 infection, the predominant CD4+ T-cell response has been associated with mild disease and enhanced early virus clearance in acute infection while its absence was associated with fatal COVID-19 outcome (4,8,9). Although at a lower level of magnitude, SARS-CoV-2 specific CD8+ in coordination with CD4+ T-cell response in acute infection seems to be essential for a good prognosis (4). Opposite to these findings, a higher magnitude and broader overall T-cell response (10, 11) and higher antibody levels against SARS-CoV-2 (12) have anti-intuitively been associated, according to the original antigenic sin theory (13), with poor disease outcome. Despite all these findings, the information about the quality and polyfunctionality of T-cell response in acute infection is scarce. The detailed and comprehensive analysis by intracellular staining (ICS) may clarify existing paradoxes about the role of T-cell response in acute infection and may provide additional immune correlates of protection.

Equally important for immune protection and recurrent COVID-19 is to analyze the immune memory after SARS-CoV-2 infection. The longevity of CD4+ T-cell and memory B cell response against the spike protein (S) seems to be stable, while CD8+ T-cell response lowered by half at six to eight months after infection (14). Moreover, it is very important to know whether the disease severity during acute infection may dictate the quality and the magnitude of long-term immune memory. Patients with post-acute symptoms showed a trend to decline IFN-γ production in N-specific CD8+ T-cells four months after infection (11). However, a detailed analysis of the quality of T-cell response at the longer term after infection is lacking. These analyses may have important implications on current vaccination strategies.

The immune memory response is not always triggered by a previous contact with SARS-CoV-2. Heterologous response in unexposed healthy donors has been found due to the sequence homology between common cold coronaviruses (HCoV) and SARS-CoV-2 (8,15–17). A detailed analysis of the correlation and the qualities of these responses is needed in order to know potential correlates of protection and vaccine responses.

In the present study, using an integrative approach combining antibody levels, SARS-CoV-2 specific CD4+ and CD8+ T-cell response, we found specific magnitude and polyfunctionality features of this response associated with disease severity in acute infection, with long-term immune memory in previously hospitalized and non-hospitalized patients and also associated to heterologous response to endemic coronaviruses.

## MATERIAL AND METHODS

### Study participants

Seventy participants with confirmed detection of SARS-CoV-2 by reverse-transcription polymerase chain reaction (RT-PCR) as previously described (18) were included. Out of these 70, 37 were hospitalized in acute phase of COVID-19 from March 25^th^ to May 8^th^ 2020, while 33 participants were recruited seven months after being diagnosed with COVID-19, from September 9^th^ to November 26^th^ 2020. These participants came from the COVID-19 patients’ Cohort Virgen del Rocio University Hospital, Seville (Spain) and the COVID-19 Cohort IIS Galicia Sur (CohVID GS), Vigo (Spain) (19). Thirty-three healthy donors (HD), pre-COVID-19 cryopreserved samples (May 12^th^ to July 18^th^ 2014) were included in the HD cohort, collected at Laboratory of HIV infection, Andalusian Health Public System Biobank, Seville (Spain) (C330024) (19). Written or oral informed consent was obtained from all participants. The study was approved by the Ethics Committee of the Virgen Macarena and Virgen del Rocio University Hospital (protocol code “pDCOVID”; internal code 0896-N-20). Hospitalized participants during the acute phase of infection were divided in Mild (n=18) or Severe (n=19), based on the highest level of disease severity during course of hospitalization. Severe participants were those who required Intensive Care Unit admission, or having ≥6 points in the score on ordinal scale (20) or death. The remaining acutely infected individuals by SARS-CoV-2 were considered mild. Blood samples were collected at a median of 3 days [interquartile range (IQR) 2.0 – 21.5] after hospitalization and 17 days [7.0 – 31.5] after symptoms onset (Table S1). The group of participants discharged after infection, included previously hospitalized (n=19) and previously non hospitalized subjects (n=14). The samples from these participants were collected after a median of 201 days [180.5 - 221] after hospitalization and 208 days [190 - 232] after symptoms onset (Table S1). Clinical and demographic data from both HD and infected subjects are described in Supplementary Table 1 (Table S1). Acutely SARS-CoV-2 infected patients and COVID-19 convalescent (previously hospitalized and not) participants were age and sex matched with HDs’ group (Table S1).

### Cell and plasma isolation

Peripheral blood mononuclear cells (PBMCs) from healthy donors and participants were isolated from peripheral blood samples using BD Vacutainer® CPT™ Mononuclear Cell Preparation Tubes (with Sodium Heparin) by density gradient centrifugation at the same day of blood collection. Afterwards, PBMCs were cryopreserved in freezing medium (90% of fetal bovine serum (FBS) + 10% dimethyl sulfoxide (DMSO)) in liquid nitrogen until further use. Plasma samples were obtained using BD Vacutainer™ PET EDTA centrifugation tubes and were cryopreserved at −80°C until further use.

### Cell stimulation

PBMCs were thawed, washed and rested for 1 h in 0.25 µL/mL DNase I (Roche Diagnostics, Indianapolis, IN)-containing R-10 complete medium (RPMI 1640 supplemented with 10% FBS, 100 U/ml penicillin G, 100 l/ml streptomycin sulfate, and 1.7 mM sodium L-glutamine). 1.5 × 10^6^ PBMCs were stimulated *in vitro* for 6 h with overlapping peptides of protein S (PepMix™ SARS-CoV-2; Spike Glycoprotein, from JPT, Berlin, Germany), 1.5 × 10^6^ with N (PepMix™ SARS-CoV-2; Nucleocapside Protein, from JPT, Berlin, Germany) and 1.5 × 10^6^ with protein S of an optimized peptide pool of endemic coronavirus (SE) (21). 1.5 × 10^6^ PBMCs incubated with the proportional amount of dimethyl sulfoxide (DMSO) were included in each batch of experiments as a negative control. The stimulation was performed in the presence of 10 µg/mL of brefeldin A (Sigma Chemical Co, St. Louis, MO) and 0.7 µg/mL of monensin (BD Biosciences) protein transport inhibitors, anti-CD107a-BV650 (clone H4A3; BD Biosciences, USA) monoclonal antibody and purified CD28 and CD49d as previously described (22). T-cell specific response was defined as the frequency of cells expressing intracellular cytokines and/or degranulation markers after stimulation with S, N and SE peptides, normalized with the unstimulated condition (background subtraction).

### Immunophenotyping and intracellular cytokine staining

Both cultured PBMCs and cells for phenotypical analysis were washed (1800 rpm, 5 min, room temperature) with Phosphate-buffered saline (PBS) and incubated 35 min at room temperature (RT) with LIVE/DEAD Fixable Aqua Dead Cell Stain (Life Technologies), anti-CD14-BV510 (clone MφP9), anti-CD19-BV510 (clone SJ25C1), anti-CD56-BV510 (clone NMCAM16.2), anti-CD3-BV711 (clone SP34-2), anti-CD45RA-FITC (clone L48), anti-CD8-APC (clone SK-1), anti-CD27-APCH7 (clone M-T271), anti-PD-1-BV786 (CD279, clone EH12-1), anti-CD38 (clone HIT2), anti-CD28 (clone CD28.2) (all of them from BD Bioscience); anti-TIGIT-PerCPCy5.5 (clone A15153G) and anti-HLA-DR (clone L243) (from BioLegend). PBMCs were washed with PBS and fixed and permeabilized with BD Cytofix/CytoPerm following manufacturer’s protocol (Cat. No. 554714, BD Bioscience), and intracellularly stained at 4°C for 30 min with anti-IL-2-BV421 (clone MQ1-17H12), anti-IFN-γ-PE-Cy7 (clone B27) (BD Bioscience), anti-TNF-α-AF700 (clone Mab11) (BD Pharmingen), anti-Perforin-PE (clone B-D48) (BioLegend). T cells were gated based on the CD3 and CD8 expression. Each subset (Total Memory, MEM; Central Memory, CM; Effector Memory, EM; and terminally differentiated effector memory, TEMRA) was gated based on CD45RA and CD27 expression (for gating strategy see Fig. S12). The specific T-cell response to each stimuli was determined by the sum of the expression of each cytokine (IFN-γ, IL-2 and TNF-α) in the different T-cell subsets. To classify an individual as a responder, this value must be higher than 0.05 (22). Flow cytometry analyses were performed on an LRS Fortessa flow cytometer using FACS Diva software (BD Biosciences). For this analysis, at least 1×10^6^ events were acquired per sample and a median of 4.72 × 10^5^ live T-cells were gated. Data were analyzed using the FlowJo 10.7.1 software (Treestar, Ashland, OR).

### Cytokine quantification

Cytokine levels were assayed in plasma samples using three different kits. sCD25 were measured by Human CD25/IL-2R alpha Quantikine ELISA Kit (R&D System, Cat# DR2A00) and IP-10 by Human IP-10 ELISA Kit (CXCL10) (Abcam, Cat# ab173194). In order to quantify IL-6, IL-8, IL-1β, TNF-α, IFN-γ, MIP-1α, MIP-1β, MILLIPLEX MAP Human High Sensitivity T Cell Panel (Merck Cat# HSTCMAG-28SK) were used. All of these kits were utilized according to the manufacturer’s instructions.

### Quantification of anti-S SARS-CoV-2 and endemic coronaviruses IgG antibodies

Anti-S IgG SARS-CoV-2 and endemic coronaviruses (NL63, OC43, 229E and HKU1) levels were measured by ELISA as previously described (4,16,23,24). Briefly, Nunc Maxisorp flat-bottomed 96-well plates (ThermoFisher Scientific #3690) were coated with 1μg/mL of recombinant SARS-CoV-2 (Sino Biological, #40589-V08B1), NL63 (Sino Biological, #40604-V08B), OC43 (Sino Biological, #40607-V08B), 229E (Sino Biological, #40605-V08B) and HKU1 (Sino Biological, #40606-V08B) Spike protein, overnight at 4°C. The following day, plates were blocked with 3% milk in phosphate buffered saline (PBS) containing 0.05% Tween-20 for 120 min at RT. Plasma samples were heat inactivated at 56°C for 45 min. Plasma was diluted 1:50 or 1:100 in 1% milk containing 0.05% Tween-20 in PBS and incubated for 90 min at room temperature. Plates were washed 4 times with 0.05% PBS-Tween-20. Secondary antibodies, streptavidin-horseradish peroxidase-conjugated mouse anti-human IgG (Hybridoma Reagent Laboratory, Baltimore, MD, #HP6043-HRP) was used at a 1: 2,000 dilutions in 1% milk containing 0.05% Tween-20 in PBS. Plates were washed 4 times with 0.05% PBS-Tween-20. The plates were developed using fast o-phenylenediamine Peroxidase Substrate (Merck, #P9187), the reaction was stopped using 1M HCl, and the optical density at 490 nm (OD490) was read on a Multiskan GO Microplate Spectrophotometer (ThermoFisher Scientific) within 2 hrs.

### Statistical Analysis

Non-parametric statistical analyses were performed using Statistical Package for the Social Sciences software (SPSS 25.0; SPSS, Inc., Chicago, IL), RStudio Version 1.3.959 and GraphPad Prism version 8.0 (GraphPad Software, Inc.). Polyfunctionality was defined as the percentage of lymphocytes producing combinations of cytokines (IL-2, TNF-α and IFN-γ), the degranulation marker CD107a and perforin. The simultaneous expression of the 3 cytokines, were also named as(3 functions, plus CD107a and/or PRF, as4 and 5 functions, respectively.. Polyfunctionality pie charts were constructed using Pestle version 1.6.2 and Spice version 6.0 (provided by M. Roederer, NIH, Bethesda, MD) and was quantified with the polyfunctionality index algorithm (25) employing the 0.1.2 beta version of the FunkyCells Boolean Dataminer software provided by Martin Larson (INSERM U1135, Paris, France). Median and interquartile ranges were used to describe continuous variables and percentages to describe categorical variables. ROUT method was utilized to identify and discard outliers. Differences between different groups were analyzed by two-tailed Mann-Whitney U test. The Wilcoxon test was used to analyze paired samples. Categorical variables were compared using the χ2 test or the Fisher’s exact test. The Spearman test was used to analyze correlations between variables. All differences with a P value of < 0.05 were considered statistically significant.

## RESULTS

### Hospitalized patients with acute SARS-CoV-2 infection show an altered T-cell phenotypic profile

Patients hospitalized with acute SARS-CoV-2 infection showed higher CD4+ and lower CD8+ T-cells levels compared with sex- and age-matched pre-COVID-19 healthy donors (HD), which resulted in higher CD4:CD8 T-cell ratio (Fig. 1A, left panel). SARS-CoV-2 infection was also associated with lower effector memory (EM) and terminally differentiated effector memory (TEMRA) CD4+ T-cell levels (Fig. 1A, middle panel), while no differences were observed in CD8+ T-cell subset levels (Fig. S1A). CD4:CD8 TEMRA ratio was lower in acute COVID-19 patients compared with HD (Fig. 1A, right panel). Analyses of T-cell activation by HLA-DR and CD38 co-expression revealed higher levels in all of CD8+ T-cell subsets and TEMRA CD4+ T-cells in SARS-CoV-2 infected patients (Fig. 1B). This was also observed for CD38 single expression in all CD8+ T-cell subsets (Fig. S1B) but not for HLA-DR single expression (Fig. S1C-D). The levels of senescent CD4+ (CD57+CD28-), but not CD8+ T-cell subsets were lower in acute infection (Fig. 1C). However, T-cell exhaustion, assayed by PD-1 and TIGIT expression and co-expression of both markers, was higher in acute SARS-CoV-2 infection in most of the T-cell subsets (Fig. 1D-F).

**Figure 1.**
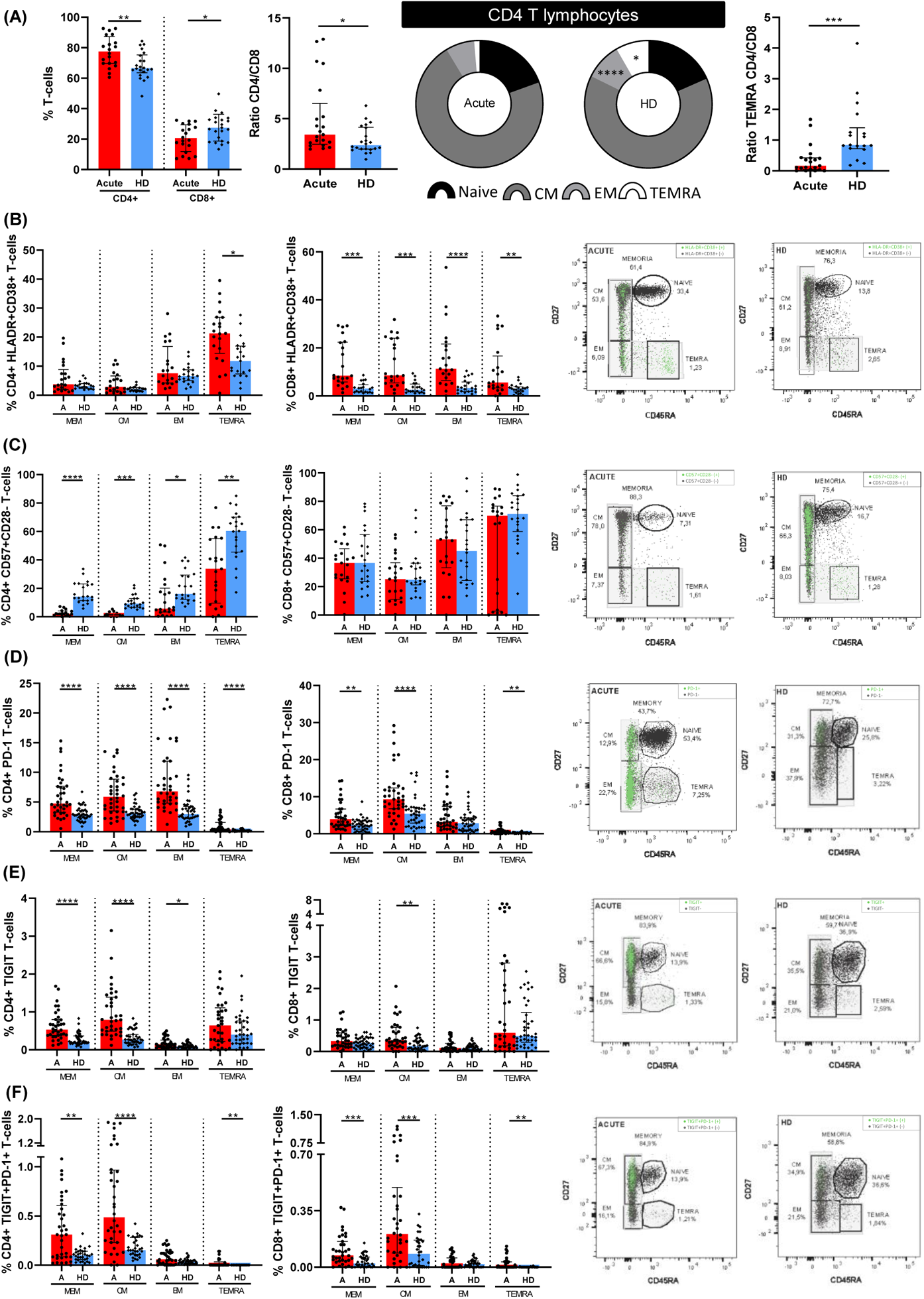
Altered CD4+ T-lymphocyte maturation phenotype and markers of T-cell activation, senescence and exhaustion in patients with acute SARS-CoV-2 infection. (A) Bar graphs representing the percentage of total CD4+ and CD8+ T cells (left panel); ratio between CD4+ and CD8+ (middle panel); and, CD4:CD8 ratio in TEMRA T-cell subset (right panel). Pie graphs show medians of each CD4+ T-cell subset in acute SARS-CoV-2 infected individual (Acute) and healthy donor (HD) groups. Each subsets from both groups were compared. Next bar graphs together with the representative dot-plots of each group mentioned above show the expression of each biomarker: (B) HLA-DR+CD38+ (activation marker); (C) CD57+CD28- (immune senescence marker); (D) PD-1+; (E) TIGIT+ and (F) PD-1+TIGIT+ T-cells (exhaustion markers). The medians with the interquartile ranges are shown. For dot-plots, green points are positive events of each biomarker, while the negatives are represented in black. *p < 0.05, **p < 0.01, ***p < 0.001, ****p < 0.0001. Mann-Whitney U test was used for groups’ comparisons and Spearman test for non-parametric correlations. Categorical variables were compared using the χ2 test or the Fisher’s exact test.

### Characteristics of SARS-CoV-2 specific T-cell response in acute hospitalized patients and healthy donors

We assayed SARS-CoV-2 specific T-cell response by intracellular cytokine staining (ICS), this technique is a well-stablished method for evaluating virus specific T-cell response (22, 26). ICS, despite of using a high number of cells, allowed us to have higher sensitivity to assay the magnitude and quality of the T-cell response. We performed CD4+ and CD8+ T-cell response specific to spike (S) and nucleocapsid (N) peptide pools. The specific T-cell response to each stimuli was determined by the sum of the expression of each assayed cytokine (IFN-γ, IL-2 and TNF-α). To classify an individual as a responder, we consider a threshold higher than 0.05%, as previously published (22). First, as expected, we observed a higher magnitude of the response in most of T-cell subsets for both peptide pools (N and S) in hospitalized acute SARS-CoV-2 infected patients (Acute) compared to HD samples (Fig. 2A-B, top panels). However, there were differences neither in the magnitude of the response nor in the proportion of responders in the TEMRA subset for both CD4+ and CD8+ T-cells and for S and N stimuli (Fig. 2A-B). In fact, there were no differences in the levels of responders for all CD8+ T-cell subsets for N peptides (Fig. 2B, bottom panels). Overall, 75% and 82% of HD had SARS-CoV-2 specific CD4+ and CD8+ T-cell response, respectively, considering S+N peptides and all T-cell subsets (Fig. 2C-D).

**Figure 2.**
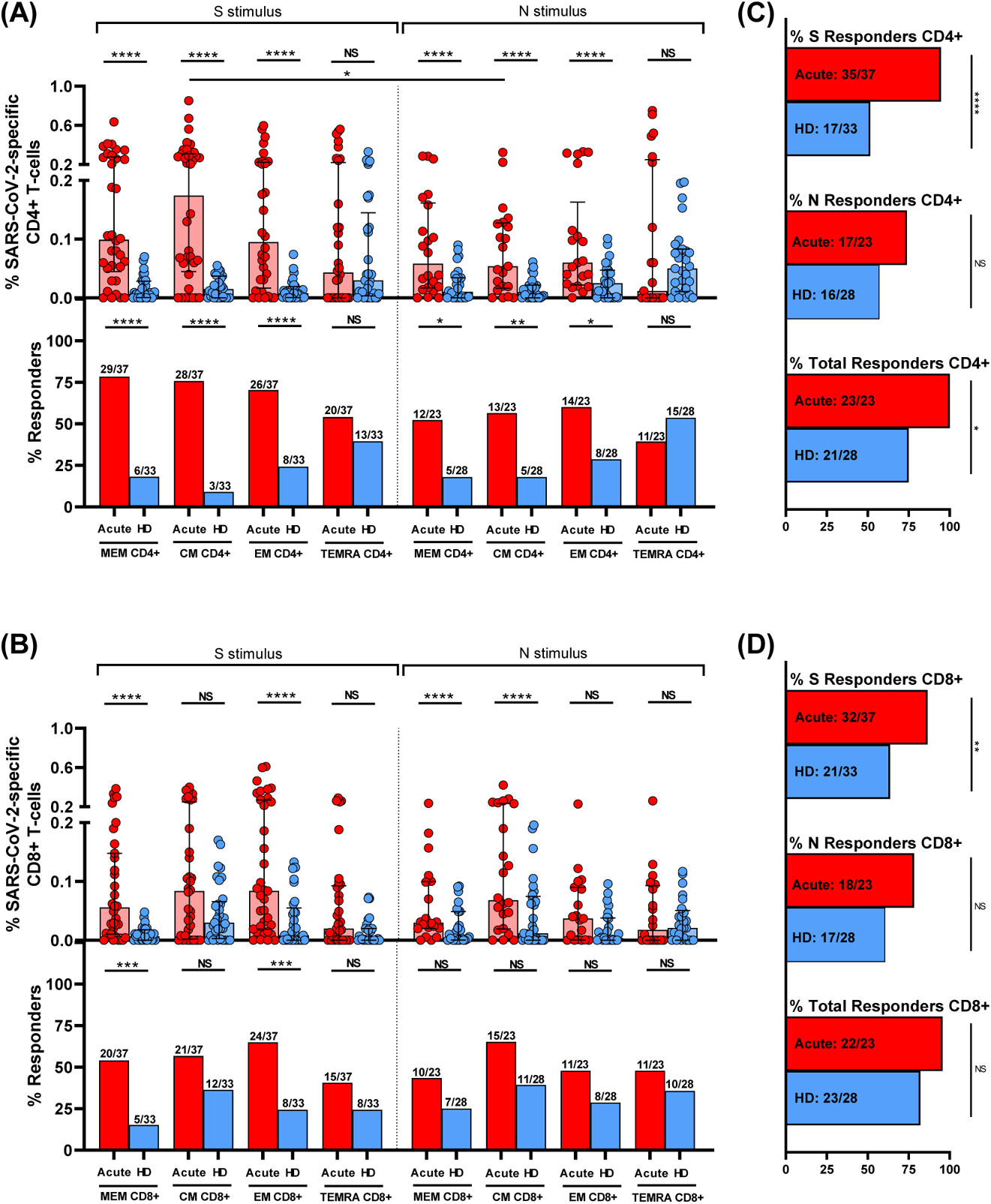
S and N specific CD4+ and CD8+ T-cell response in acute SARS-CoV-2 infection individuals and healthy donors. **(A)** and **(B)** Bar graphs in top panels represent percentage of S and N specific CD4+ and CD8+ T-cell response in SARS-CoV-2 infected patients (red) and healthy donors (blue) (upper panels). Bar graphs in low panels also show the number and percentage of responders, considering a responder subjects as those with the percentage of SARS-CoV-2-specific T-cells higher than 0.05% considering the sum of IFN-γ, TNF-α and IL-2 production. **(C)** and **(D)** Bar graphs describe the number and percentage of responders for S peptide pool, as the sum of any CD3+CD4+ or CD3+CD8+ T-cell subset (%S Responders); for N peptide pool, as the sum of any CD3+CD4+ or CD3+CD8+ T-cell subset (%N Responders) and the total responders as the sum of CD3+CD4+ or CD3+CD8+ S and N responses (% of total Responders). The medians with the interquartile ranges are shown. Each dot represents an individual. *p < 0.05, **p < 0.01, ***p < 0.001, ****p < 0.0001. Mann-Whitney U test was used for groups’ comparisons. Categorical variables were compared using the χ2 test or the Fisher’s exact test.

Second, comparing the response to S and N peptide pools, there was a higher magnitude of response to S compared to N stimulus in central memory (CM) CD4+ T-cell (p=0.042) and a trend in total memory (MEM) CD4+ T-cells (p=0.126) (Fig. 2A, top panels), however there were no differences for CD8+ T-cell subsets (Fig. 2B, top panels). Additionally, a cumulative SARS-CoV-2-specific T-cell measurement was calculated as the sum of the S and N responses (Fig. S2). Our data show that, all the patients had detectable SARS-CoV-2 specific T-cell response considering together the response against S and N peptide pools and to all the CD4+ and CD8+ T-cell subsets (Fig.2C-D, Fig. S2).

Finally, when comparing CD4+ and CD8+ T-cell response in hospitalized patients (Acute), the magnitude of SARS-CoV-2 specific total memory (MEM) CD4+ T-cell response (Fig. 2A, top panel) was higher compared MEM CD8+ T-cell response (Fig. 2B, top panel) for protein S (p=0.048), but not different for the rest of subsets and for protein N (Fig. 2A-B, top panels). In the same way, there was a higher percentage of responders for MEM CD4+ T-cells compared to MEM CD8+ T-cells in S protein (p=0.025), there were no differences in the proportion of responders for the rest of subsets for S and N proteins (Fig. 2A-B, bottom panels).

### IFN-γ and IL-2 polyfunctional response in S-specific CD4+ T-cells are differentially associated with disease severity while IL-2 production in S-specific CD8+ T-cells is associated with mild disease

We next analyzed in acute infection the association of S-specific CD4+ T-cell response with disease severity in hospitalized patients segregated as mild and severe patients. The S-specific CD4+ T cell response was significantly higher in the TEMRA CD4+ T-cell subset in severe compared to mild patients (Fig. 3A). When individual cytokine production was analyzed, these higher levels were attributed to IFN-γ production in S-specific TEMRA CD4+ subset (Fig. 3B; Fig S3A), but not for IL2 or TNF-α (Fig. S3B). Multiple combination of cytokines, together with CD107a and perforin expression revealed that combinations only including IFN-γ+ CM (Fig. 3C, Fig. S3C) and TEMRA cells (Fig. S3D) were increased in severe compared with mild patients. The same occurred for combinations including IFN-γ+ and TNF-α+ CM cells (Fig. S3E). However, combinations including IL-2, such as IL2+TNF-α+ MEM cells were increased in mild compared to severe patients (Fig. 3D). In fact, S-specific MEM CD4+ T-cell polyfunctionality was higher in mild patients, mainly because of increased bi-functional combinations including IL-2 (IL2/TNF-α, IL2/IFN-γ and IL-2/perforin) that were not present in severe patients, where IFN-γ/TNF-α combination was predominant (Fig. 3E). It is also important to highlight that perforin expression was higher in MEM and EM subsets of mild patients in comparison with severe patients (Fig. S3F). This was reflected in a higher S-specific MEM polyfunctional index in mild compared to severe patients (Fig. 3F). Additionally, we observed that the bulk of S-specific CD4+ T-cell response in the different subsets was inversely associated with different inflammatory markers, while specific combinations including IFN-γ were directly associated with plasmatic IP-10 levels (Fig. S4). Overall, a high polyfunctional S-specific CD4+ T-cell response biased to IL-2 production was associated with mild disease, while combinations only including IFN-γ were associated with severe disease outcomes.

**Figure 3.**
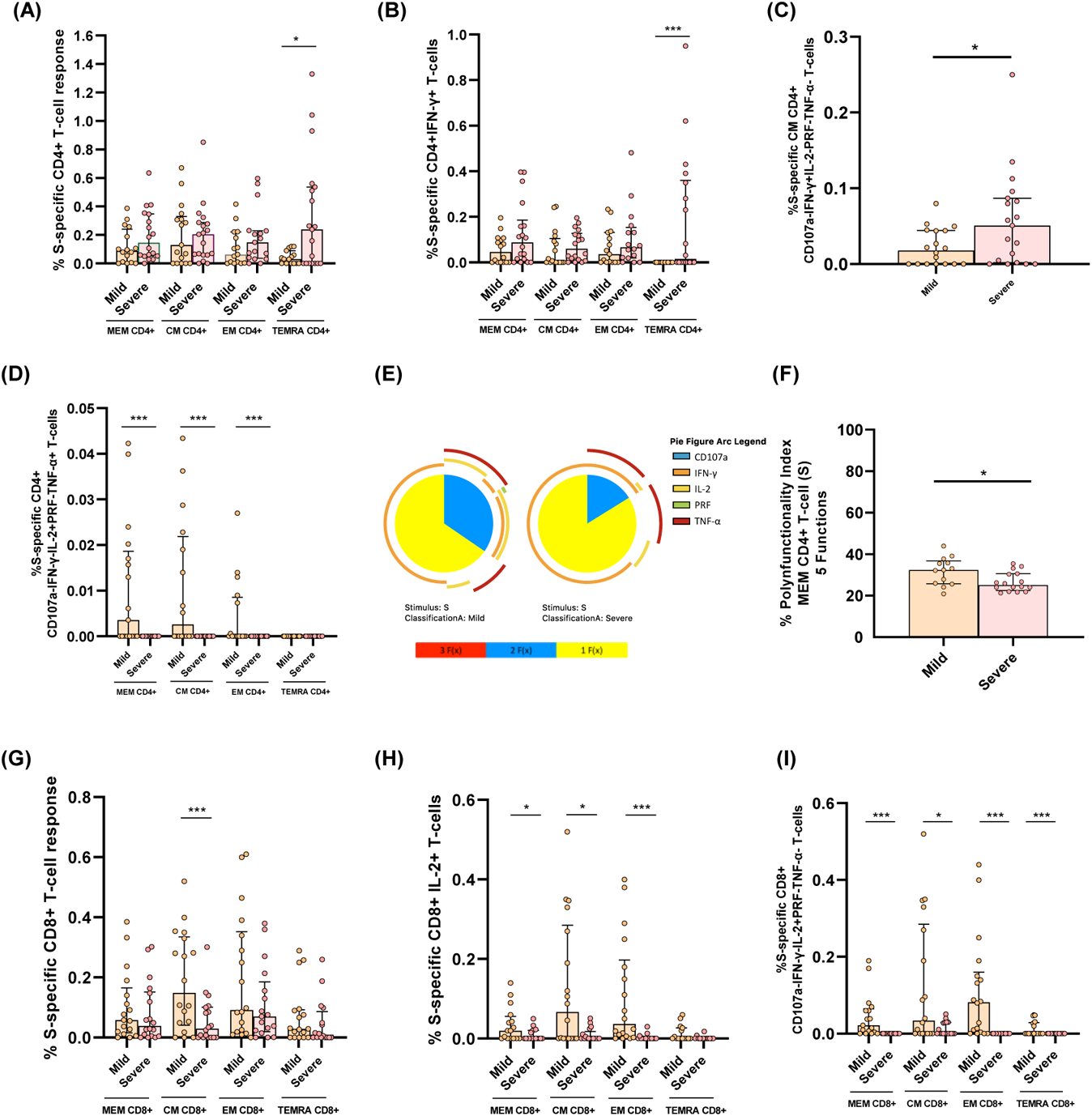
S-specific CD4+ and CD8+ T-cell response is associated with disease severity in acute SARS-CoV-2 infection. **(A)** Bar graphs show S-specific CD4+ T-cell response, considering the sum of IFN-γ, TNF-α and IL-2 production, in the different CD4+ T-cell subsets, in mild and severe acute patients’ groups. **(B)** S-specific CD4+ T-cell response considering the levels of cells producing IFN-γ. **(C)** S-specific CM CD4+ T-cell levels of combinations only including IFN-γ+ cells for five (IFN-γ, TNF-α, IL-2, CD107a and PRF) functions. **(D)** S-specific CD4+ T-cell levels in the different T-cell subsets of combinations including IL-2+ and TNF-α+ cells for five (IFN-γ, TNF-α, IL-2, CD107a and PRF) functions. **(E)** S-specific MEM CD4 T-cell polyfunctionality pie charts. Each sector represents the proportion of S-specific CD4 T-cells producing two (blue) and one (yellow) function. Arcs represents the type of function (IFN-γ, TNF-α, IL-2, CD107a and PRF) expressed in each sector. Permutation test, following the Spice version 6.0 software was used to assess differences between pie charts. **(F)** Polyfunctional index bar graph of S-specific MEM CD4+ polyfunctionality, for five functions. **(G)** Bar graphs show S-specific CD8+ T-cell response, considering the sum of IFN-γ, TNF-α and IL-2 production, in the different CD8+ T-cell subsets, in mild and severe acute patients’ groups, **(H)** S-specific CD8+ T-cell response considering the levels of cells producing IL-2 and **(I)** S-specific CD8+ T-cell levels of combinations only including IL-2+ cells for five (IFN-γ, TNF-α, IL-2, CD107a and PRF) functions. The medians with the interquartile ranges are shown. Each dot represents a patient. *p < 0.05, **p < 0.01, ***p < 0.001. Mann Whitney U test was used for groups’ comparisons.

In relation to S-specific CD8+ T-cells, the bulk of CM CD8+ T-cell response was higher in mild compared to severe patients (Fig. 3G). We observed that the cytokine responsible of these differences was IL-2, which presented higher levels in MEM, CM and EM S-specific CD8+ T-cells in mild subjects (Fig. 3H; Fig. S5A) while very low levels and no differences were observed in IFN-γ+ and TNF-α+ production (Fig. S5B). These results were confirmed by combinations only including IL-2 without the expression of the rest of the cytokines, CD107a and perforin, in the same subsets: MEM, CM and EM (Fig. 3I). Similar results were observed for three and four functions (Fig. S5C). In summary, IL-2 production in not terminally differentiated S-specific CD8+ T-cells was associated with mild disease progression in hospitalized acute SARS-CoV-2 infected patients.

### Polyfunctional N-specific CD4+ T-cell response is associated with mild disease in acute SARS-CoV-2 hospitalized patients

We also analyzed in detail the quality of N-specific T-cell response. MEM and CM IL2+ and EM TNF-α+ N-specific CD4+ T cell levels were higher in mild compared to severe patients (Fig. 4A-B). We did not observe differences for the bulk of IFN-γ+ N-specific CD4+ T-cell response (Fig. S6A-B). Following the same profile of S-specific CD4+ T-cell response, combinations including only IL2+ and TNF-α+ in MEM, CM and EM N-specific CD4+ T-cells were associated with mild disease progression (Fig. 4C). Besides, we observed higher levels of combinations with triple cytokine positive MEM, CM and EM CD4+ T-cells in mild compared to severe patients (Fig. 4D). In fact, MEM N-specific response showed a higher proportion of triple and a variety of double combinations (Fig. 4E), likewise a higher MEM polyfunctional index in mild compared to severe patients (Fig. 4F). These results were reproduced in polyfunctionality of CM and EM subsets with three and four functions that were also associated with mild disease progression (Fig. S6C). We did not observe great differences in N-specific CD8+ T-cell response according with disease severity, only higher levels of combinations with only IFN-γ+ T-cells in severe compared to mild patients (Fig. S6D).

**Figure 4.**
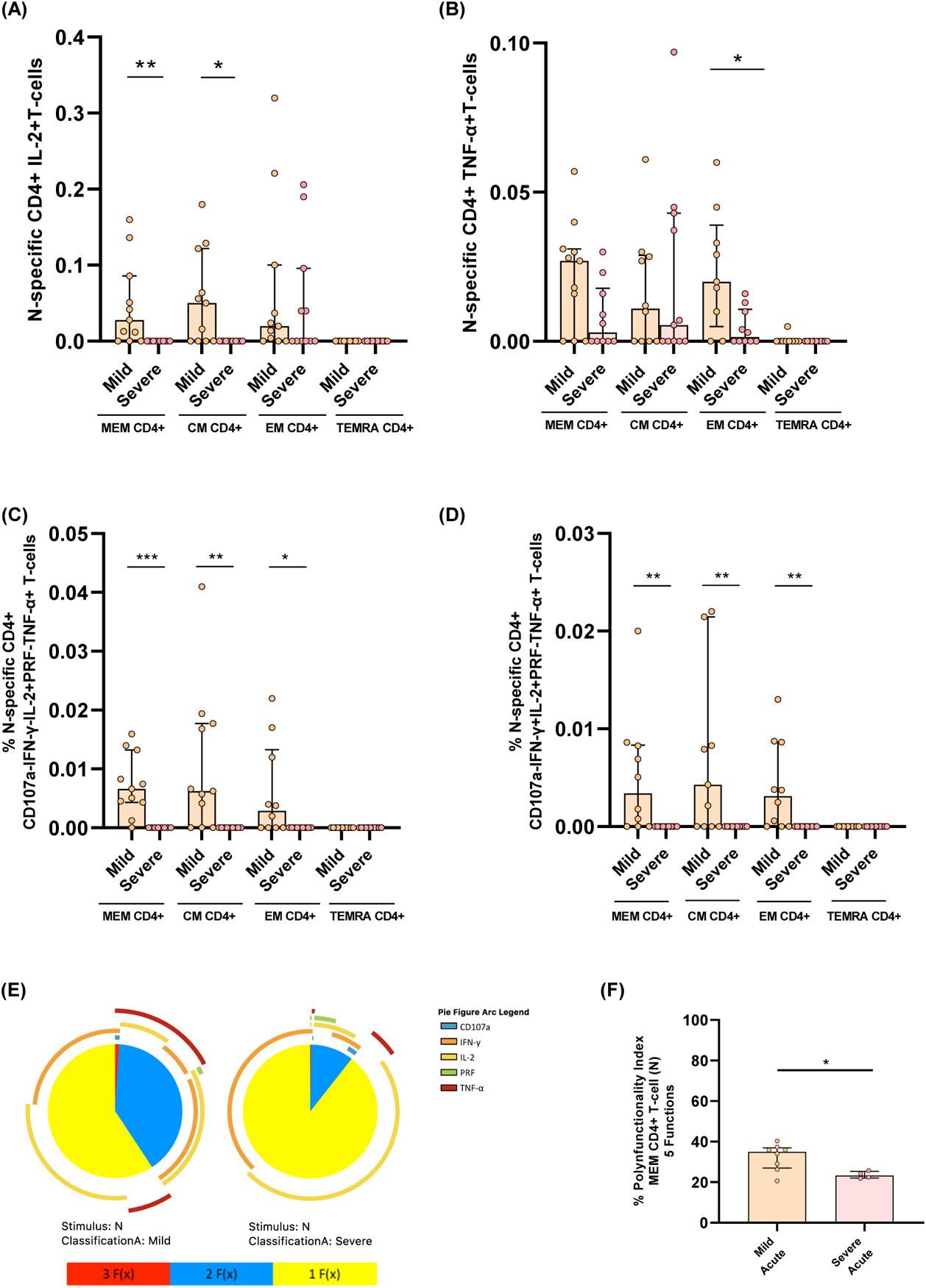
Cytokine combinations and polyfunctional N-specific CD4+ T-cell response are associated with COVID-19 progression. **(A)** N-specific CD4+ T-cell response considering the levels of cells producing IL-2 in the different CD4+ T-cell subsets, in mild and severe acute patients’ groups. **(B)** N-specific CD4+ T-cell response considering the levels of cells producing TNF-α in the different CD4+ T-cell subsets, in mild and severe acute patients’ groups. **(C)** N-specific CD4+ T-cell levels in the different T-cell subsets of combinations including IL-2+ and TNF-α+ cells for five (IFN-γ, TNF-α, IL-2, CD107a and PRF) functions. **(D)** N-specific CD4+ T-cell levels in the different T-cell subsets of combinations including IL-2+, TNF-α+ and IFN-γ+ T-cells for five (IFN-γ, TNF-α, IL-2, CD107a and PRF) functions. **(E)** N-specific MEM CD4 T-cell polyfunctionality pie charts. Each sector represents the proportion of N-specific CD4 T-cells producing three (red), two (blue) and one (yellow) function. Arcs represents the type of function (IFN-γ, TNF-α, IL-2, CD107a and PRF) expressed in each sector. Permutation test, following the Spice version 5.2 software was used to assess differences between pie charts. **(F)** Polyfunctional index bar graph of N-specific MEM CD4+ polyfunctionality, for five functions. The medians with the interquartile ranges are shown. Each dot represents a patient. *p < 0.05, **p < 0.01, ***p < 0.001. Mann-Whitney U test was used for groups’ comparisons.

### Similar magnitude of SARS-CoV-2 specific T-cell response in previously hospitalized and non-hospitalized patients seven months after infection

In addition to the analyses in the acute phase, we analyzed the magnitude of SARS-CoV-2 specific T-cell response seven months after SARS-CoV-2 infection in two group of individuals: i) those previously hospitalized during acute infection and ii) without previous hospitalization. First, we analyzed CD4+ T-cell response. No differences were observed in the magnitude of N- and S-specific T-cell response, except for higher S-specific TEMRA CD4+ T-cell levels in non-hospitalized patients compared to previously hospitalized patients (Fig. 5A, top panel). Despite the 18% lower expression of TEMRA CD4+ T-cells in previously hospitalized responders, this difference was not statistically significant (Fig. 5A, bottom panel). No differences in the proportion of responders was observed in the rest of subsets and neither for S- nor N-protein (Fig. 5A, bottom panel). Considering the cumulative SARSCoV-2-specific CD4+ T-cell response (sum of the S and N response) (Fig. S7A) in all the T-cell subsets, all patients, with the exception of one in the group of previously hospitalized patients, had detectable CD4+ T-cell response (Fig. 5B). We also compared the magnitude of S- versus N-specific CD4+ T-cell response in both groups. We found that the magnitude of MEM and CM response was higher in S compared to N in both, previously hospitalized (p=0.008; p=0.008, respectively) and non-hospitalized patients (p=0.014; p=0.009, respectively) seven months after SARS-CoV-2 infection (Fig. 5A, top panel). Secondly, we analyzed the magnitude of SARS-CoV-2 specific CD8+ T-cell response and we found that previously hospitalized patients presented higher S-specific EM T-cells compared to non-hospitalized patients (Fig. 5C, top panel). There were no differences for the rest of subsets or stimuli between both groups (Fig. 5C, top panel). We also did not find differences in the percentage of responders (Fig. 5C, bottom panel). The analysis of the cumulative SARS-CoV-2-specific CD8+ T-cell response (Fig. S7B), showed that 88.9% and 92.8% for previously hospitalized and non-hospitalized, respectively, had detectable SARS-CoV-2-specific CD8+ T-cell response considering all T-cell subsets and S or N stimuli (Fig. 5D). Furthermore, we found that the magnitude of MEM and EM response was higher in N compared to S peptides in non-hospitalized patients (p=0.026; p=0.024, respectively) while no differences were found in previously hospitalized patients seven months after SARS-CoV-2 infection (Fig. 5C). Finally, we compared the magnitude of response between CD4+ and CD8+ T-cells. We found that in both groups, MEM and CM S-specific CD4+ T-cell response was higher than in CD8+ T-cells (p=0.005 and p=0.036 for previously hospitalized; p=0.004 and p=0.013 for non-hospitalized, respectively), while no differences were found for N stimulus (Fig. 5A-C).

**Figure 5.**
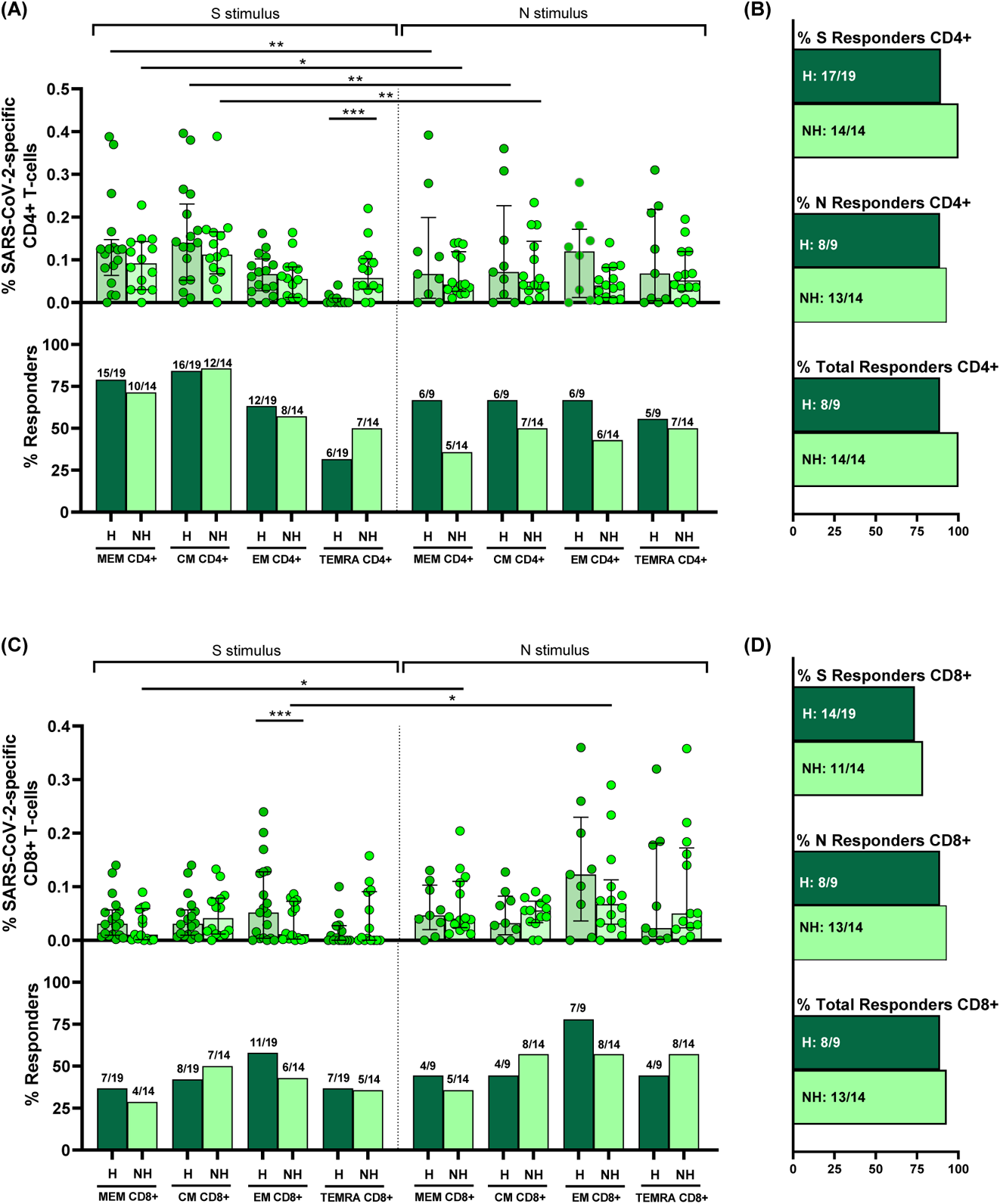
S and N specific CD4+ and CD8+ T-cell response are present in previously hospitalized (H) and non-hospitalized (NH) patients seven months after SARS-CoV-2 infection. **(A)** and **(C)** Bar graphs represent percentage of S and N specific CD4+ and CD8+ T-cell response in previously hospitalized (H) (dark green) and non-hospitalized (NH) (light green) subjects (top panels). Bar graphs also show the number and percentage of responders, considering a responder subjects as those with the percentage of SARS-CoV-2-specific T-cells higher than 0.05% considering the sum of IFN-γ, TNF-α and IL-2 production (bottom panels). **(B)** and **(D)** Bar graphs describe the number and percentage of responders for S peptide pool, as the sum of any CD3+CD4+ or CD3+CD8+ T-cell subset (%S Responders); for N peptide pool, as the sum of any CD3+CD4+ or CD3+CD8+ T-cell subset (%N Responders) and the total responders as the sum of CD3+CD4+ or CD3+CD8+ S and N responses (% of total Responders). The medians with the interquartile ranges are shown. Each dot represents an individual. *p < 0.05, **p < 0.01, ***p < 0.001, ****p < 0.0001. Mann-Whitney U test was used for groups’ comparisons. Categorical variables were compared using the χ2 test or the Fisher’s exact test.

### Higher CD4+TIGIT+ T-cell and differential quality of S-specific CD4+ CM T-cell levels in previously hospitalized compared to non-hospitalized patients seven months after infection

After finding a similar magnitude of SARS-CoV-2 specific T-cell response in both groups of individuals seven months after infection, we assayed the quality of T-cell response and exhaustion markers in previously hospitalized and non-hospitalized patients. The TIGIT expression in all the CD4+ T-cell subsets were higher in previously hospitalized than in non-hospitalized patients (Fig. 6A). We did not find differences between groups in TIGIT+ CD8+ T-cells (Fig. S8A). Likewise, PD-1 expression was similar in all T-cell subsets in both groups (Fig. S8B-C). When we analyze multiple combination of cytokines, previously hospitalized patients showed higher levels of S-specific CM CD4+ T-cell with combinations only including TNF-α (Fig. 6B, left panel; Fig. S8D). Furthermore, S-specific CM CD4+ T-cell response was more polyfunctional in non-hospitalized patients compared with those previously hospitalized (Fig. 6B, right panel). In the same line, N-specific T-cell response also contained higher levels of combinations only including IFN-γ for EM CD4+ T-cells and only including TNF-α for CM CD8+ T-cells in previously hospitalized than in non-hospitalized patients (Fig. 6C-D; Fig. S8E-F). However, non-previously hospitalized convalescent patients showed N-specific CM and EM CD8+ T-cells with higher production of IL-2 (Fig. 6E-D, right panel) and perforin (Fig. 6F), respectively.

**Figure 6.**
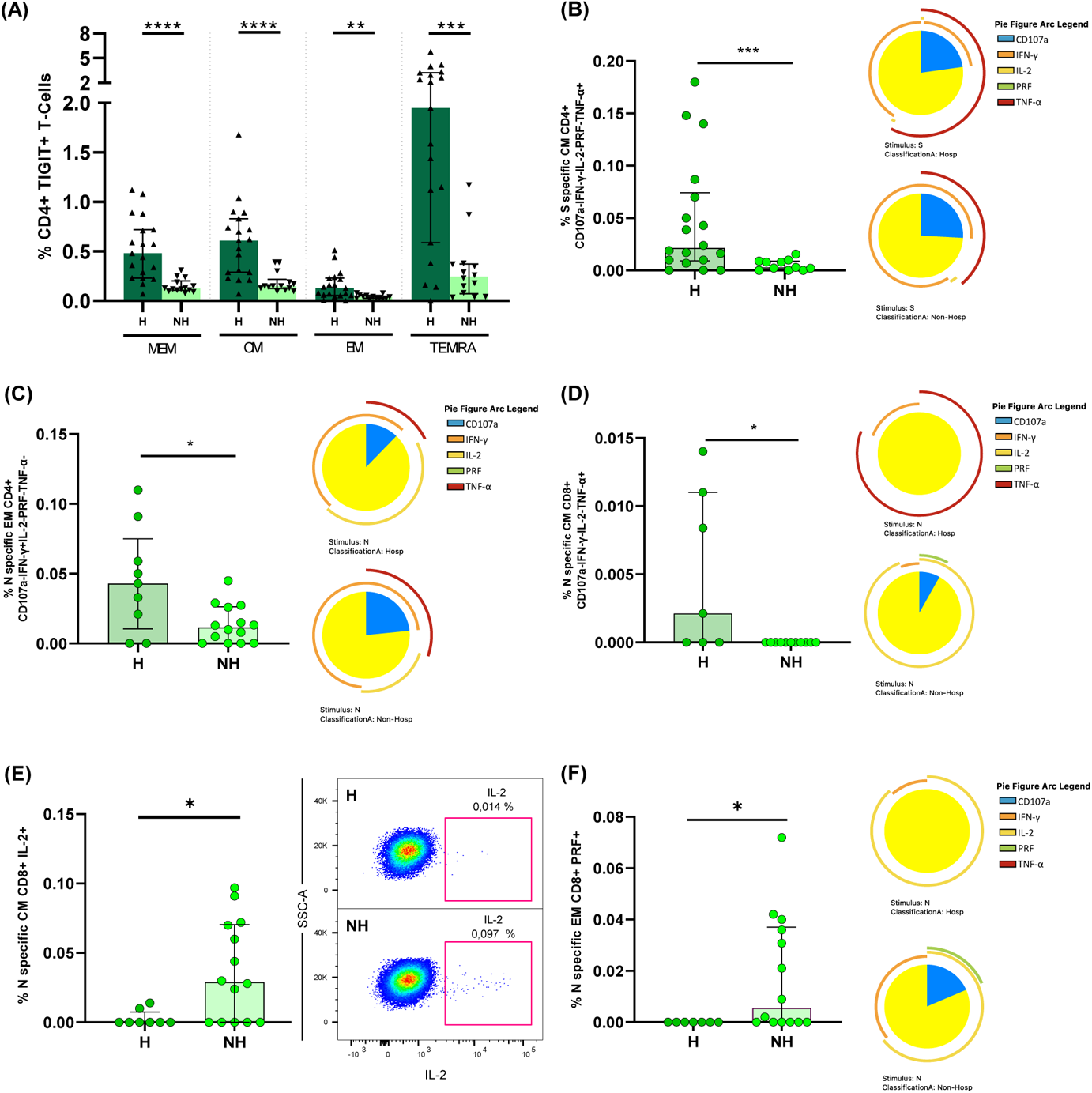
Previously hospitalized subjects during acute infection showed higher TIGIT+ CD4+ T-cell levels and lower polyfunctional S- and N-specific T-cell response than non-hospitalized subjects seven months after SARS-CoV-2 infection. **(A)** TIGIT expression in each CD4+ T-cell subset in previously hospitalized and non-hospitalized subjects seven months after SARS-CoV-2 infection. **(B)** S-specific CM CD4+ T-cell levels of combinations only including TNF-α+ cells for five (IFN-γ, TNF-α, IL-2, CD107a and PRF) functions (left panel). S-specific CM CD4 T-cell polyfunctionality pie charts (right panel). **(C)** N-specific EM CD4+ T-cell levels of combinations only including IFN-γ+ cells for five (IFN-γ, TNF-α, IL-2, CD107a and PRF) functions (left panel). N-specific EM CD4 T-cell polyfunctionality pie charts (right panel). **(D)** N-specific CM CD8+ T-cell levels of combinations only including TNF-α+ cells for five (IFN-γ, TNF-α, IL-2, CD107a and PRF) functions (left panel). N-specific CM CD8 T-cell polyfunctionality pie charts (right panel). **(E)** N-specific CM CD8+ T-cell levels of cells producing IL-2 (left panel). Representative dot plot showing IL-2 production in N-specific CM CD8+ T-cells (right panel). **(F)** N-specific EM CD8+ T-cell levels of cells producing PRF (left panel). N-specific EM CD8 T-cell polyfunctionality pie charts (right panel). For all the pie charts each sector represents the proportion of SARS-CoV-2-specific T-cells producing two (blue) and one (yellow) function. Arcs represents the type of function (IFN-γ, TNF-α, IL-2, CD107a and PRF) expressed in each sector. Permutation test, following the Spice version 6.0 software was used to assess differences between pie charts. Each dot represents an individual. *p < 0.05, **p < 0.01, ***p < 0.001, ****p < 0.0001. Mann-Whitney U test was used for groups’ comparisons.

### Anti-S IgG levels are associated with disease severity and differentially with SARS-CoV-2-specific T-cell response in acute and convalescent subjects seven months after infection

Next, we analyzed antibody levels against S protein and the association of this humoral response with disease severity and T-cell immunity. In acute infection, we observed a trend to increased antibody levels in severe compared to mild patients (Fig. 7A). Seven months after SARS-CoV-2 infection anti-S IgG levels remained high, similar to severe patients in acute infection and at higher levels compared to mild patients in both previously hospitalized and in non-hospitalized patients (Fig. 7A). As expected, all the groups had higher antibody levels compared to HD (Fig. 7A). In relation to T-cell response, in general, SARS-CoV-2 specific T-cell response was inversely associated with anti-S IgG levels in acute infection (Fig. S9A), while a direct correlation was observed seven months after infection (Fig. S9B). A representative example was the inverse correlation of S-specific EM CD4+ T-cell producing IL2 in acute infection (Fig. 7B) compared to the direct correlation found seven months after infection (Fig. 7C).

**Figure 7.**
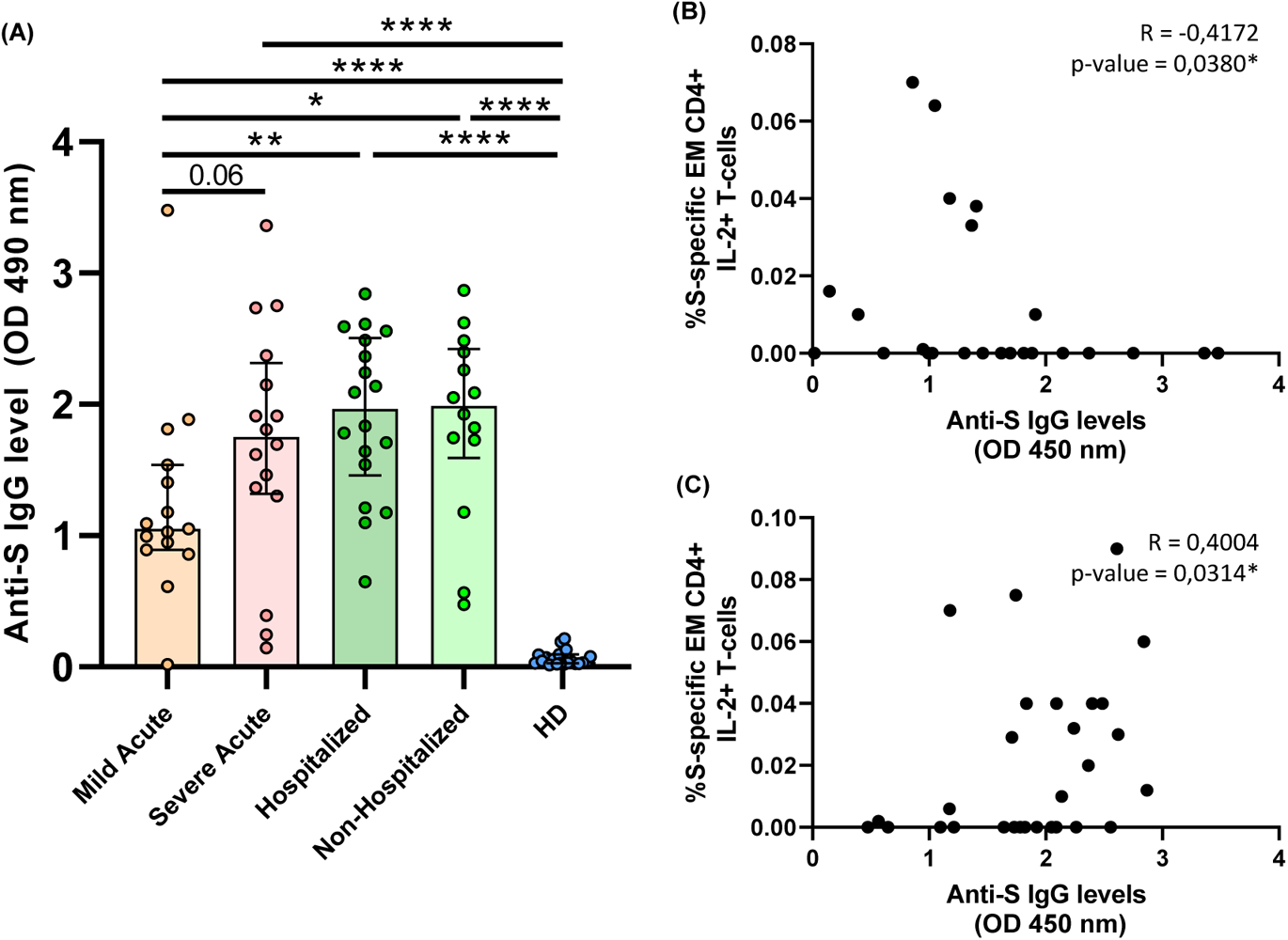
Anti-S IgG levels are associated with COVID-19 severity and correlate with S-specific T-cell response. **(A)** Bar graphs represent anti-S IgG levels in each study group. **(B)** Correlation graphs between anti-S IgG levels and the percentage of S-specific EM CD4+ IL2+ T-cell response in acute infection. **(C)** Correlation graphs between anti-S IgG levels and the percentage of S-specific EM CD4+ IL2+ T-cell response previously hospitalized patients seven months after SARS-CoV-2 infection. Each dot represents an individual. *p < 0.05, **p < 0.01, ***p < 0.001, ****p < 0.0001. Mann-Whitney U test was used for groups’ comparisons and Spearman test for non-parametric correlations.

### Heterologous SARS-CoV-2 response is associated with the magnitude and the quality of endemic coronavirus response

Similar to previously reported (21), we found that a high percentage of HD (pre-COVID-19 samples) presented detectable CD4+ and CD8+ SARS-CoV-2 specific T-cell response (75% and 82%, respectively) (Fig. 2). In order to characterize this immune response, we performed anti-S IgG levels and specific T-cell response by ICS using an optimized peptide pool for the four human endemic coronaviruses (21). In acute SARS-CoV-2 infected participants we observed a direct correlation of anti-S SARS-CoV-2 IgG levels with those of three out of the four endemic coronaviruses (HCoV-NL63, -OC43 and -HKU1) (Fig. S10). When we split this group, we only found a positive correlation of anti-S SARS-CoV-2 IgG and anti-S HCoV-NL63 and -OC43 levels in severe (Fig. S10) but no correlation was found in mild patients (Fig. S10). In HD we also found a positive correlation of anti-S SARS-CoV-2 IgG and anti-S HCoV-OC43, -229E and HKU-1 levels (Fig. S10). Finally, in all the groups together, anti-S SARS-CoV-2 IgG levels were directly associated with anti-S IgG levels of the beta-coronaviruses HCoV-OC43 and -HKU1 (Fig. S10). After that, we performed S-specific T-cell response to the optimized peptide pool of endemic coronaviruses (SE) in HD. We found detectable SE T-cell response in all the CD4+ and CD8+ T-cell subsets (Fig. 8A). Analyzing the bulk of SE T-cells reported higher levels of response in the TEMRA and CM subsets, in CD4+ and CD8+ T-cells, respectively. Overall, the proportion of responders was 55.6% for CD4+ and 72% for CD8+ T-cells and 80% the global response (Fig. 8A). The response to human endemic coronaviruses correlated to S-specific for SARS-CoV-2 in CM and EM CD4+subsets (Fig. 8B-C) and in CM CD8+ subset (Fig. 8D). Attending to the quality of this response, it was mainly monofunctional and IL-2 production prevailed in CD4+ and CD8+ T-cells respect to other cytokines (Fig. 8E; Fig. S11A). Interestingly, combinations including IL-2, but not IFN-γ, in response to human endemic coronaviruses correlated with S-specific SARS-CoV-2 response, for CD4+ MEM (Fig. 8F), CM and EM subsets (Fig. S11B-C) and CD8+ CM (Fig. 8G).

**Figure 8.**
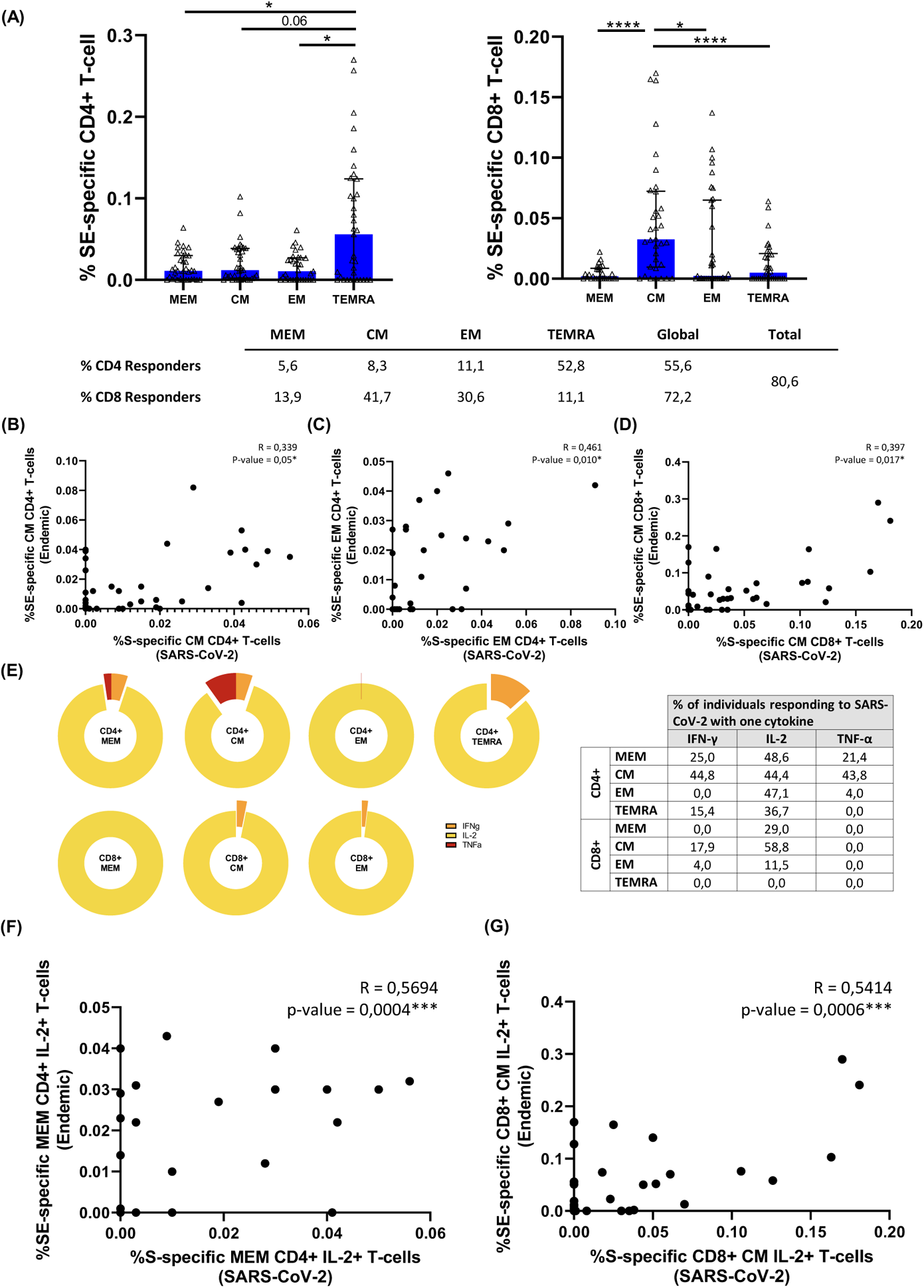
Characteristics of the cross-reactive T-cell response quality between SARS-CoV-2 and endemic coronaviruses. **(A)** Bar graph represent the SE-specific T-cell response in each CD4+ (left panel) and CD8+ (right panel) T-cell subsets. The table shows the percentage of responders considering a responder subjects as those with the percentage of SE-specific T-cells higher than 0.05% considering the sum of IFN-γ, TNF-α and IL-2 production. **(B)** Correlation between S-Specific and SE-specific CM CD4+ T cell levels. **(C)** Correlation between S-Specific and SE-specific EM CD4+ T cell levels and **(D)** Correlation between S-Specific and SE-specific CM CD8+ T-cell levels in healthy donors. **(E)** Pie graphs represent IFN-γ, IL-2 and TNF-α expression in each T-cell subset where median percentages of this expression are shown in right table. **(F)** Correlation between S-Specific and SE-specific MEM CD4+ IL-2+ T cells and **(G)** S-Specific and SE-specific CM CD8+ IL-2+ T cell levels. Each dot represents an individual. *p < 0.05, **p < 0.01, ***p < 0.001, ****p < 0.0001. Mann-Whitney U test was used for groups’ comparisons and Spearman test for non-parametric correlations.

## DISCUSSION

In the present study, analyzing 103 subjects, we describe features of SARS-CoV-2 specific humoral and T-cell response differentially associated with disease severity in hospitalized patients during acute infection. This response is long-lasting seven months after infection independently whether patients were previously hospitalized or not, although previous hospitalization was associated with exhausting T-cell features present in acute infection. Finally, we comprehensively analyzed the features of the high levels of cross-reactive response between SARS-CoV-2 and human endemic coronaviruses in healthy donors.

We used ICS for the systematic analysis of SARS-CoV-2 specific T-cell response. ICS is a technique commonly used for analyzing T-cell response against viral infections (22, 26) and can be complementary to other strategies as T-cell receptor dependent activation induced marker (AIM) (16,27,28). Although a high amount of cells is needed, a comprehensive cytokine dependent functional characterization of virus specific T-cell response can be achieved (5). We analyzed the response against protein S and N, because these are the main targets, in terms of magnitude, of SARS-CoV-2 specific T-cell response (16).

Firstly, we found that all patients in acute infection, independently of disease severity, had detectable SARS-CoV-2-specific T-cell response, as a summation of S+N response and considering all CD4+ and CD8+ T-cell subsets. These data were remarkable based on the high activation and inhibitory receptor T-cell levels found in the present study in response to SARS-CoV-2 infection. Patients in acute infection showed low CD8+ T-cell levels and consequently high CD4+:CD8+ T-cell ratio compared to HD together with high T-cell inhibitory receptor levels, such as high levels of PD-1, as previously reported (9, 29), and TIGIT, likewise high levels of activation in all the CD8+ T-cell subsets but only for terminally differentiated CD4+ T-cells. This scenario was compatible with a prominent T-cell migration to damage tissue (30, 31) which was associated with lower TEMRA CD4+:CD8+ T-cell ratio, because of high levels of TEMRA CD4+HLA-DR+CD38+ T-cells accompanied by low peripheral CD4+ T-cell senescent levels, pointing out to preferential tissue recruitment of these cells.

According with previous studies, we found higher levels of response in CD4+ compared to CD8+ T-cells and against S than N protein in CD4+ T-cells (29). This was confirmed in the MEM subset. The higher CD4+ compared to CD8+ T-cell response levels may be due to the use of optimized peptide pools for MHC-II. However, higher CD4+ compared to CD8+ T cell response levels have been traditionally associated with control of SARS-CoV-1 infection (6, 32). In our cohort, the high number of responders could be mainly due to CD8+ T-cells (96%) while other cohorts only found 53% of responders in this subset (4),. In this sense, the use of ICS for assaying T-cell response may achieve a higher sensitivity compared to other methods.

In acute infection, some studies have associated a deleterious effect of SARS-CoV-2-specific T-cell response with disease progression (10, 11) while others have shown a beneficial role associated with mild disease in acute hospitalized patients (9). Results presented herein may clarify this paradox. Severe compared to mild patients showed higher IFN-γ but lower IL-2+TNF-α+ S- and N-specific CD4+ T-cell levels. These results means that studies using only IFN-γ ELISPOT technology (10, 33) would consider SARS-CoV-2-specific T-cell response in acute infection as deleterious. However, we found that combinations including IL-2 were polyfunctional, including the cytotoxic marker perforin, pointing out to a higher antiviral activity with a classical signature of canonical Th1 cells (4,5,8,10) associated with mild disease in hospitalized patients. This fact was supported by S-specific CD8+ T-cell response, which mainly consisted in IL-2 production, in this case in monofunction, at higher levels in mild patients. Besides, it is important to highlight the higher polyfunctionality found in N compared to S protein. Polyfunctional combinations of three functions (IFN-γ+IL-2+TNF-α+) were at higher levels in MEM, CM and EM CD4+ T-cells in mild compared to severe patients in response to N protein. A successful outcome of acute disease may come for the combination and coordination of CD4+, CD8+ T-cell response and antibody production against SARS-CoV-2 (4). For having an integrated picture of acute anti-SARS-CoV-2 response, we also assayed anti-S IgG levels that in accordance with previous studies (9, 12), were at higher levels in severe patients. Although overall, in acute infection, as previously reported (9, 10), S-specific T-cell response was directly associated with anti-S IgG levels, we found that the production of IL-2 by EM S-specific T-cells was inversely associate with antibody production. In the same line, overall S-specific CD4+ T-cell response was inversely associated with inflammatory markers; however, combinations including IFN-γ were directly associated with IP-10 plasmatic levels which has previously associated with disease progression (34, 35). Altogether, these results define two different quality profiles of humoral and S/N-specific T-cell response associated with diseases progression in hospitalized patients: i) a mild disease progression profile associated with IL-2 production, inversely correlated with anti-S IgG levels and associated with a higher T-cell polyfunctionality, which should promote CD4+ T-cell proliferation and CD4+ T-cell help to CD8+ T-cells together with antiviral potential enabling rapid virus clearance; and ii) a severe disease progression profile consisting in high anti-S IgG levels and combinations only including IFN-γ, mainly in terminally differentiated T-cells with absence of perforin production, no CD8+ T-cell help and limited antiviral potential what may favor the failure to early control of the virus and poor disease outcome.

Next, we sought to analyze the immune memory to SARS-CoV-2 seven months after infection in two groups of subjects with different course of the disease: patients that overcame the disease without the need of hospitalization and previously hospitalized patients. We observed that in both groups, all subjects displayed detectable T-cell response, considering S+N response and all CD4+ and CD8+ T-cell subsets. This is in accordance with immune memory found to SARS-CoV infection, which have been shown to last for years (36) and agreed with the magnitude of T-cell response found eight months after SARS-CoV-2 infection in previously non-hospitalized subjects (14). Although in that study using AIM they found only SARS-CoV-2 Specific CD8+ T-cell response in 50% while we observed 91% of responders but 75% in previously hospitalized patients (14). Despite the general absence of difference in the magnitude of T-cell response between both groups, in terms of quality of this response, previously hospitalized patients showed higher T-cell exhaustion levels (TIGIT and PD-1 expression) and higher S and N-specific T-cell levels of combinations of only including IFN-γ and TNF-α production compared to non-hospitalized patients. Additionally, non-hospitalized patients presented higher IL-2 and perforin production in N-specific CD8+ T-cells compatible with a preserved antiviral activity. This profile is reminiscent of the one found in severe compared to mild patients in acute infection. However, on the contrary to what happened in acute disease, seven months after infection in previously hospitalized subjects, anti-S IgG levels were directly, not inversely, associated with SARS-CoV-2 specific T-cell response, especially that enriched in IL-2 production which was associated with a good prognosis in acute infection. These results demonstrate that anti-SARS-CoV-2 humoral and cellular response are long-lasting and robust at least seven months after infection in both non-hospitalized and previously hospitalized patients. However, previously hospitalized patients showed T-cell exhaustion and some signs of SARS-CoV-2-specific T-cell response associated with disease progression in acute infection, although this response was more IL-2 biased, which was associated with good prognosis. This defects found in previously hospitalized patients seven months after infection may be selectively associated with long-COVID symptoms, as has been recently reported four months after infection (11); however, we cannot confirm it because information about long-lasting symptoms were not recorded in this cohort as this was not the aim of the present study.

Finally, we found high levels of cross-reactive CD4+ and CD8+ T-cell response to SARS-CoV-2 (75% and 82%, respectively) in pre-COVID-19 HD samples. These levels were even higher than those found in previous cohorts showing 20-50% of cross-reactivity (8,16,36,37). Using an optimize peptide pool (21) for the four endemic coronaviruses: NL63, OC43, 229E and HKU1, we found that endemic S-specific CD4+ and CD8+ T-cell response was directly correlated with SARS-CoV-2 S-specific CD4+ and CD8+ T-cell response in EM and CM subsets. These results confirm cross-reactive SARS-CoV-2 specific T-cell response with endemic coronavirus. Comprehensively analyses of endemic S-specific T-cell response was mainly induced by TEMRA CD4+ T-cells and CM CD8+ T-cells and we found that the response was totally biased to IL-2 production, what may explain some previously published results using IFN-γ ELISPOT that did not find cross-reactive response (10). In fact, combinations including IL-2, but not IFN-γ, were the ones associated with SARS-CoV-2 S-specific CD4+ and CD8+ T-cell response. These results suggest that this pre-existing T-cell memory may have the potential to induce low COVID-19 fatality. However, whether a different quality of endemic T-cell specific response, based mainly in IL-2 or IFN-γ production, may contribute to variations in COVID-19 progression is currently unknown.

One limitation of this study is that all patients included were recruited in the first wave of COVID-19 in Spain, in those dates, experimental therapies with very limited but transitory immunosuppressive effects were administered what may have affected the levels of immune parameters in acute infection. However, in those patients who were treated with IFN-β and corticosteroids, samples were collected time enough after these therapies to reverse the potential effects (24 days [7 – 28] and 9 days [1 – 21], respectively) what may have not affected the results presented herein. Seven months post-infection, one potential bias may come from the group of hospitalized subjects that were composed by patients with previous mild (42%) or severe (58%) disease, however, as no differences were found in any parameter associated to T-cell response (data not shown) between this two subgroups, they formed part of the same group of previously hospitalized and were compared with non-hospitalized patients. ICS needs a notable amount of cells to be assayed, this avoid us to perform endemic virus-specific T-cell response in COVID-19 samples; however, it has allowed us to obtain comprehensive data about the quality of SARS-CoV-2 specific T-cell response. Finally, anti-S IgG levels were assayed against the whole S protein and not for RBD, cross-reactive reaction cannot be excluded and results have to be interpreted taking this into account. In the same way, further research is needed to confirm and correlate our T-cell response profile with neutralizing antibody levels and B-cell polyfunctionality.

In summary, our results gain insights in the characteristics of T-cell response associated with disease severity in acute infection, supporting important information about correlates of immune protection, such as a broader polyfunctional CD4+ T-cell response with predominance of IL-2 production also present in SARS-CoV-2 specific CD8+ T-cell response, distinguished mild disease progression from severe COVID-19 characterized by an inefficient monofunctional IFN-γ+ CD4+ T-cell response in acute hospitalized patients. However, independently of previous hospitalization, SARS-CoV-2 specific T-cell response was robust seven months after infection, although some defects associated with T-cell exhaustion were observed in previously hospitalized patients.

These results have implications for protective immunity against SARS-CoV-2 and recurrent COVID-19 and may help to identify populations, apart from the classical risk ones, that are in the need of new boosting of existing vaccines, as well as for improving the design of new prototypes in order to achieve of broader long-lasting protection against COVID-19.

## ACKNOWLEDGMENTS

This study would not have been possible without the collaboration of Virgen del Rocio University Hospital COVID Team and COHVID-GS and IISGS Biobank. All of the patients, nursing staff, and data managers who have taken part in this project. We thank Prof. Shane Crotty for the review of the manuscript.

Members of COHVID-GS (Galicia Sur Health Research Institute): Alejandro Araujo, Jorge Julio Cabrera, Víctor del Campo, Manuel Crespo, Alberto Fernández, Beatriz Gil de Araujo, Carlos Gómez, Virginia Leiro, María Rebeca Longueira, Ana López-Domínguez, José Ramón Lorenzo, María Marcos, Alexandre Pérez, María Teresa Pérez, Lucia Patiño, Sonia Pérez, Silvia Pérez-Fernández, Eva Poveda, Cristina Ramos, Benito Regueiro, Cristina Retresas, Tania Rivera, Olga Souto, Isabel Taboada, Susana Teijeira, María Torres, Vanesa Val, Irene Viéitez

Members of the Virgen del Rocio University Hospital COVID Team: Jose Miguel Cisneros Herreros, César Sotomayor, Cristina Roca Oporto, Nuria Espinosa Aguilera, Luis Giménez Miranda, José Molina, Almudena Aguilera, Clara Aguilera, Teresa Aldabo-Pallas, Verónica Alfaro-Lara, Cristina Amodeo, Javier Ampuero, Maribel Asensio, Bosco Barón-Franco, Lydia Barrera-Pulido, Rafael Bellido-Alba, Máximo Bernabeu-Wittel, Claudio Bueno, Candela Caballero-Eraso, Macarena Cabrera, Enrique Calderón, Jesús Carbajal-Guerrero, Manuela Cid-Cumplido, Juan Carlos Crespo, Yael Corcia-Palomo, Elisa Cordero, Juan Delgado, Alejandro Deniz, Reginal Dusseck-Brutus, Ana Escoresca Ortega, Fatima Espinosa, Michelle Espinoza, Carmen Ferrándiz-Millón, Marta Ferrer, Teresa Ferrer, Ignacio Gallego-Texeira, Rosa Gámez-Mancera, Emilio García, María Luisa Gascón-Castillo, Aurora González-Estrada, Demetrio González, Rocío González-León, Carmen Grande-Cabrerizo, Sonia Gutiérrez, Carlos Hernández-Quiles, Concepción Herrera-Melero, Marta Herrero-Romero, Carmen Infante, Luis Jara, Carlos Jiménez-Juan, Silvia Jiménez-Jorge, Mercedes Jiménez-Sánchez, Julia Lanseros-Tenllado, José María Lomas, Álvaro López, Carmina López, Isabel López, Luis F López-Cortés, Rafael Luque-Márquez, Daniel Macías-García, Luis Martín-Villén, Aurora Morillo, Dolores Nieto-Martín, Francisco Ortega, Amelia Peña-Rodríguez, Esther Pérez, Rafaela Ríos, Jesús F Rodríguez, María Jesús Rodríguez-Hernández, Santiago Rodríguez-Suárez, Ángel Rodríguez-Villodres, Nieves Romero-Rodríguez, Ricardo Ruiz, Zaida Ruiz de Azua, Celia Salamanca, Sonia Sánchez, Javier Sánchez-Céspedes, Victor Manuel Sánchez-Montagut, Alejandro Suárez Benjumea, and Javier Toral.

## FUNDING

Consejeria de Transformacion Economica, Industria, Conocimiento y Universidades Junta de Andalucia (research Project CV20-85418) (ERM)

NIH contract 75N9301900065 (AS, DW)

Consejeria de Salud Junta de Andalucia (Research Contract RH-0037-2020 to JV)

Instituto de Salud Carlos III (CP19/00159 to AGV, FI17/00186 to MRJL, FI19/00083 to CGC, CM20/00243 to APG and COV20/00698 to support COHVID-GS)

Red Temática de Investigación Cooperativa en SIDA (RD16/0025/0020; RD16/0025/0026), which is included in the Acción Estratégica en Salud, Plan Nacional de Investigación Científica, Desarrollo e Innovación Tecnológica, 2008 to 2011 and 2013 to 2016

Instituto de Salud Carlos III, Fondos FEDER. ERM was supported by the Spanish Research Council (CSIC).

“Contratación de Personal Investigador Doctor” supported by the European Social Fund and Junta de Andalucía (PAIDI DOCTOR-Convocatoria 2019-2020). (FJO, SB).

## AUTHOR CONTRIBUTIONS

ALPG and CGC performed the experiments and curated the data. ALPG and JV analyzed and interpreted the data and participated in writing of the paper. FJO, ASG, MTR, EMM, TGP and AGV participated in data collection, data analysis and interpretation and performed experiments, MRJL, SB, MRIB participated in paper data interpretation, IRJ, MPG, MGG and MAG participated in data collection, JPS, MDNA, AFV, APG and LFLC participated in data collection and paper interpretation. DW, AS, LFLC and EP participated in paper data analysis, patient and data collection, interpretation/discussion of the results and coordination. ERM, participated in data analysis and interpretation, writing, conceived the idea and coordinate the project.

## CONFLICT OF INTERESTS

A.S. is a consultant for Gritstone, Flow Pharma, Arcturus, Immunoscape, CellCarta, OxfordImmunotech and Avalia. LJI has filed for patent protection for various aspects of T cell epitope and vaccinedesign work. All other authors declare that they have no competing financial interests.

## Notes

### Competing Interest Statement

The authors have declared no competing interest.

