## Supplementary figures for "Deciphering the quality of SARS-CoV-2 specific T-cell response associated with disease severity, immune memory and heterologous response"

Alberto Pérez-Gómez et al.

*Corresponding author information:

Ezequiel Ruiz-Mateos Carmona, Ph.D.

### SUPPLEMENTARY MATERIALS

#### Supplementary Figure 1

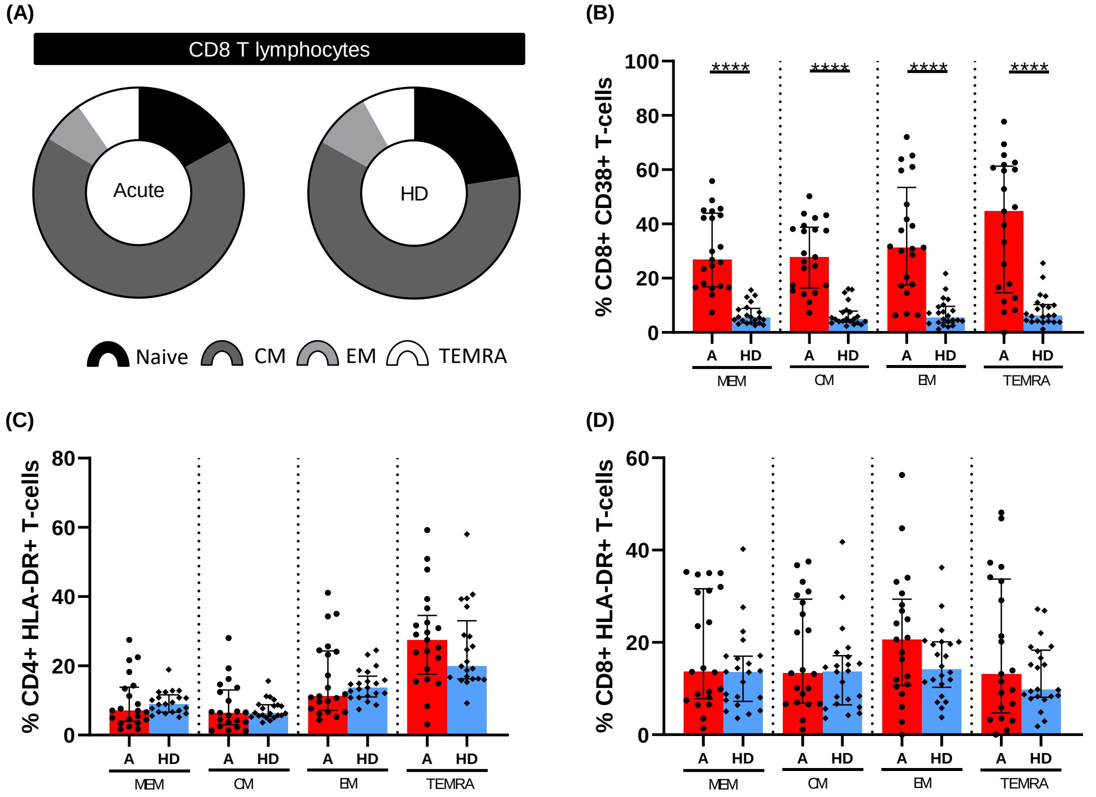


**Supplementary Figure 1. CD8+ T-cells maturation phenotype and activation markers. (A)** Pie graphs show medians of each CD8+ T-cell subset in acute SARS-CoV-2 infected individuals and healthy donor groups. Bar graphs represent the percentage of **(B)** CD8+CD38+, **(C)** CD4+HLA-DR+ and **(D)** CD8+HLA-DR+ T-cells. The medians with the interquartile ranges are shown. *$p < 0.05$, **$p < 0.01$, ***$p < 0.001$, ****$p < 0.0001$. Mann-Whitney U test was used for groups' comparisons and Spearman test for non-parametric correlations.

**Supplementary Figure 2**

**(A)**

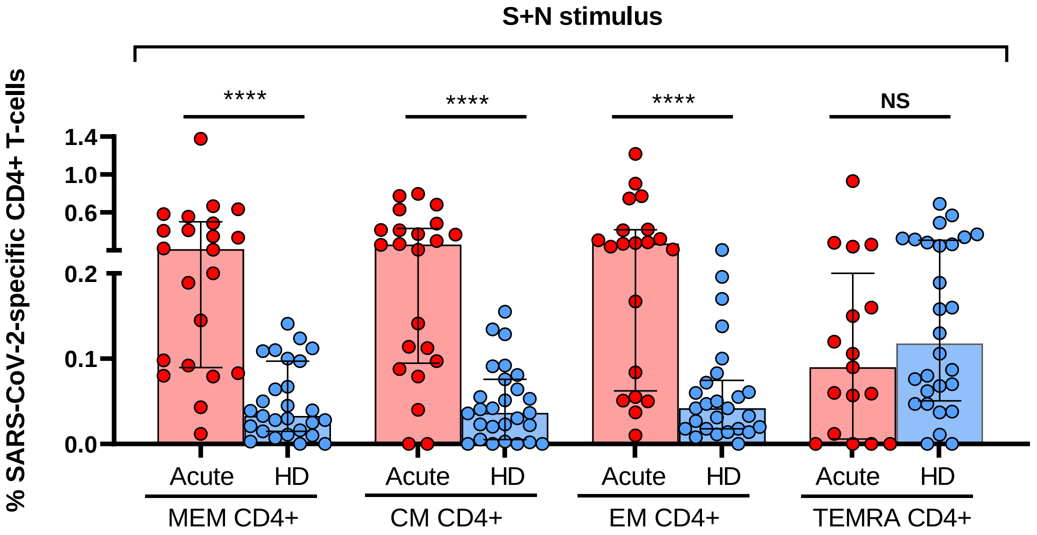


**(B)**

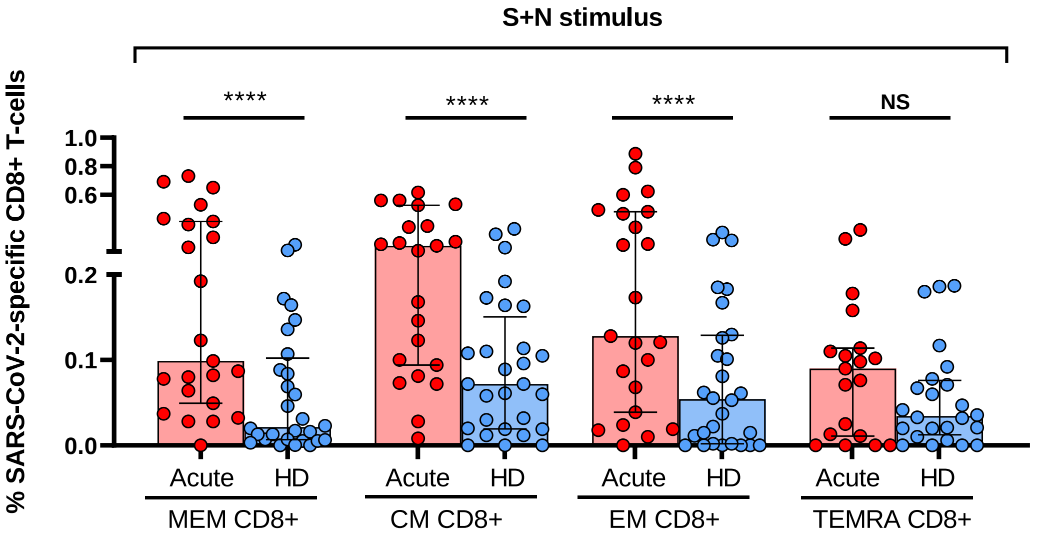


**Supplementary Figure 2. Combined SARS-CoV-2 specific T-cell response to N and S proteins in acute SARS-CoV-2 infected individuals and healthy donors. (A)** Bar graphs represent the percentage of S plus N-specific CD4+ T-cell response. **(B)** Bar graphs represent the percentage of S plus N-specific CD8+ T-cell response. The medians with the interquartile ranges are shown. Each dot represents an individual. *p < 0.05, **p < 0.01, ***p < 0.001, ****p < 0.0001. Mann-Whitney U test was used for groups' comparisons.

**Supplementary Figure 3**

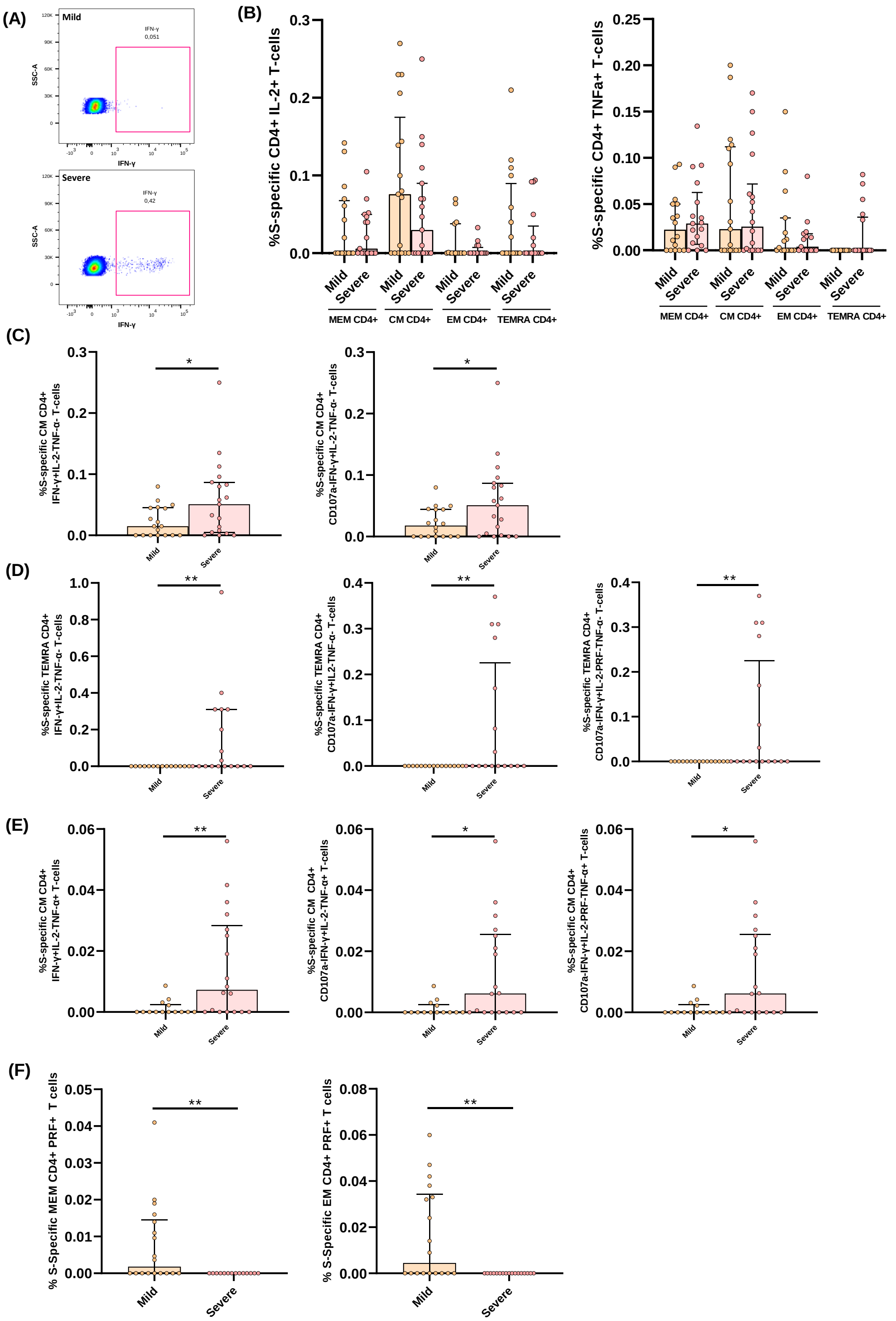


**Supplementary Figure 3. Additional S-specific CD4+ T-cell response features associated with disease severity in acute SARS-CoV-2 infection. (A)** Representative dot-plot of expression of IFN-γ+ in CD4+ TEMRA cells. **(B)** Bar graphs represent S-specific CD4+ T-cell response considering the levels of cells producing IL-2 (left panel) and TNF-α (right panel). **(C)** S-specific CM CD4+ T-cell levels of combinations only including IFN-γ+ cells for three (IFN-γ, TNF-α and IL-2) and four (IFN-γ, TNF-α, IL-2 and CD107a) functions. **(D)** S-specific TEMRA CD4+ T-cell levels of combinations only including IFN-γ+ cells for three (IFN-γ, TNF-α and IL-2), four (IFN-γ, TNF-α, IL-2 and CD107a) and five (IFN-γ, TNF-α, IL-2, CD107a and PRF) functions. **(E)** S-specific CM CD4+ T-cell levels of combinations including IFN-γ+ and TNF-α+ cells for three (IFN-γ, TNF-α and IL-2), four (IFN-γ, TNF-α, IL-2 and CD107a) and five (IFN-γ, TNF-α, IL-2, CD107a and PRF) functions. **(F)** Percentage of S-specific production of: PRF+ in CD4+ total memory (MEM) and central memory (CM) T-cell subsets. The medians with the interquartile ranges are shown. Each dot represents a patient. *$p < 0.05$, **$p < 0.01$. Mann-Whitney U test was used for groups' comparisons.

**Supplementary Figure 4**

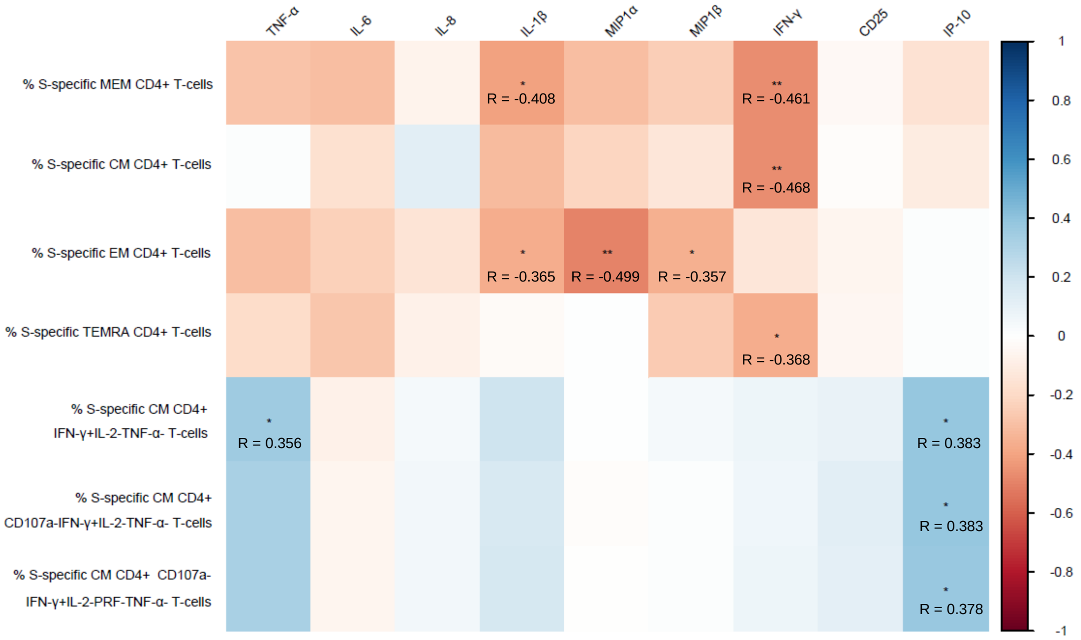


**Supplementary Figure 4. S-specific T-cell response is differentially associated to inflammatory markers in acute SARS-CoV-2 infected patients**. Correlation matrix representing associations between representative parameters of S-specific CD4+ T-cell response with inflammatory markers including TNF-α, IL-6, IL-8, IL1-β, MIP1-α, MIP1-β, IFN-γ, CD25 and IP-10, in acute SARS-CoV-2 infected patients. Blue color represents positive correlations and red color shows negative correlations. The intensity of the color indicates the R coefficient. *$p < 0.05$, **$p < 0.01$, ***$p < 0.001$. Spearman test was used for non-parametric correlations.

**Supplementary Figure 5**

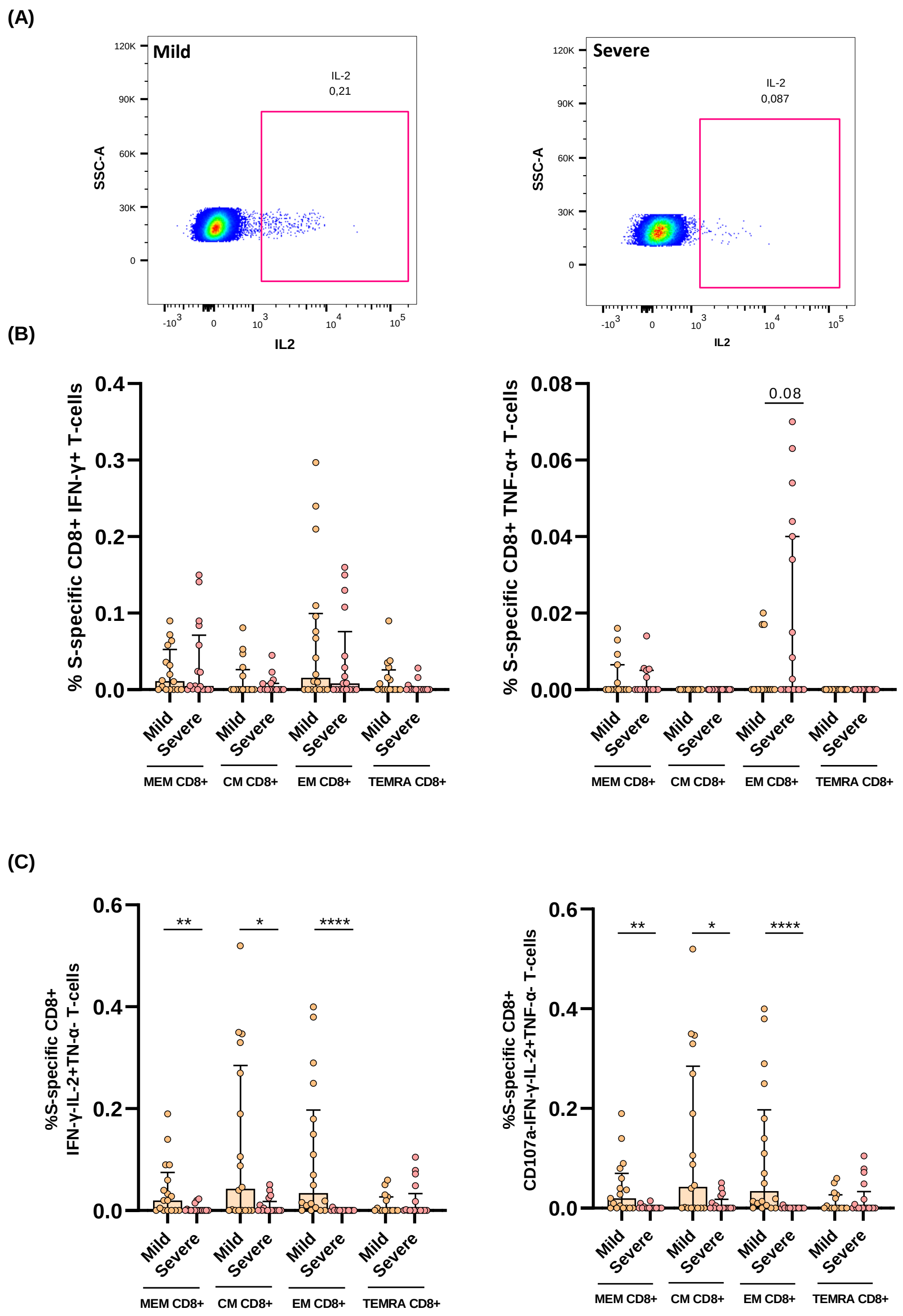


**Supplementary Figure 5. Additional features of S-specific CD8+ T-cell response associated with disease severity. (A)** Representative dot-plot of IL-2 expression in S-specific CM CD8+ T-cells. **(B)** Bar graphs represent S-specific CD8+ T-cell response considering the levels of cells producing IFN-γ (left panel) and TNF-α (right panel). **(C)** S-specific CD8+ T-cell levels of combinations only including IL-2+ cells for three (IFN-

$\gamma$, TNF-$\alpha$ and IL-2 and CD107a) and four (IFN-$\gamma$, TNF-$\alpha$, IL-2 and CD107a) functions. The medians with the interquartile ranges are shown. Each dot represents a patient. *$p < 0.05$, **$p < 0.01$, ***$p < 0.001$, ****$p < 0.0001$. Mann-Whitney U test was used for groups' comparisons.

**Supplementary Figure 6**

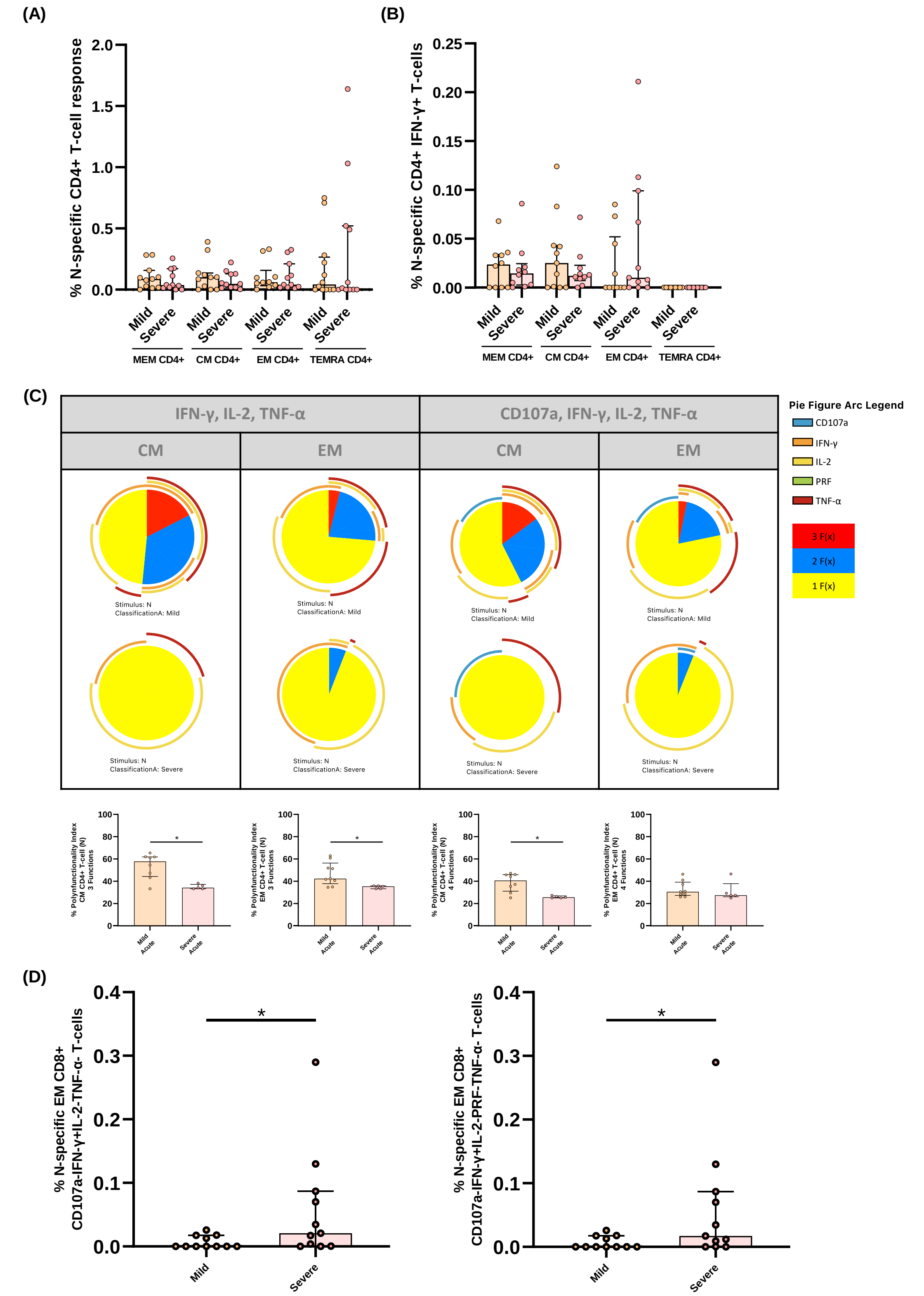


**Supplementary Figure 6. Additional features of N-specific T-cell response associated with disease progression. (A)** Bar graphs show N-specific CD4+ T-cell response, considering the sum of IFN-γ, TNF-α and IL-2 production, in the different CD4+ T-cell subsets, in mild and severe acute patients' groups, **(B)** N-specific CD4+ T-cell response considering the levels of cells producing IFN-γ. **(C)** N-specific CM and EM CD4 T-cell polyfunctionality pie charts for three and four functions. Each sector represents the proportion of N-specific CD4 T-cells producing three (red), two (blue) and one (yellow) function. Arcs represents the type of function (IFN-γ, TNF-α, IL-2, CD107a and PRF) expressed in each sector. Permutation test, following the Spice version 6.0 software was used to assess differences between pie charts. **(D)** N-specific EM CD8+ T-cell levels of combinations only including IFN-γ+ cells for four (IFN-γ, TNF-α, IL-2 and CD107a) and five (IFN-γ, TNF-α, IL-2, CD107a and PRF) functions. The medians with the interquartile ranges are shown. Each dot represents a patient. *$p < 0.05$, **$p < 0.01$, ***$p < 0.001$. Mann-Whitney U test was used for groups' comparisons.

Supplementary Figure 7

(A)

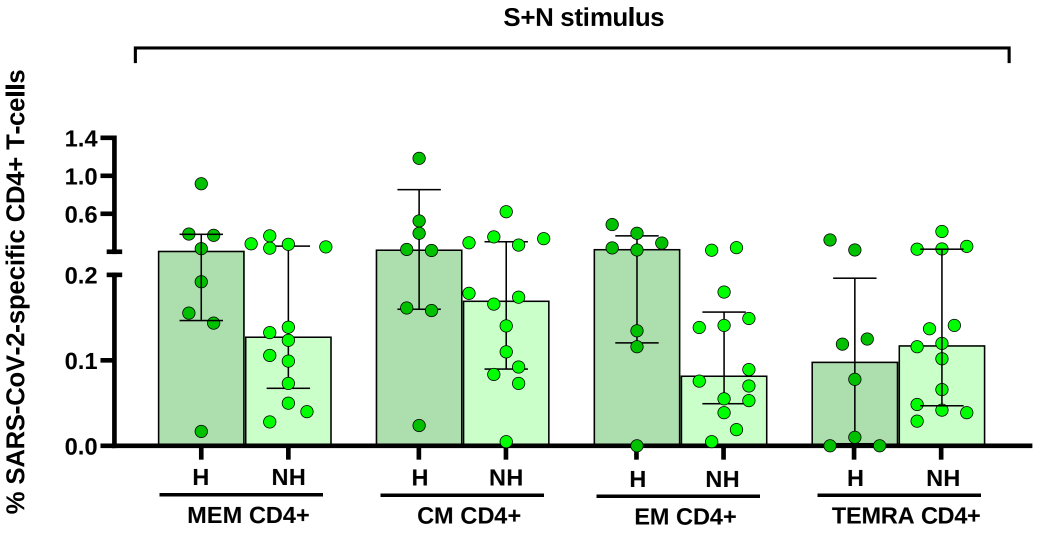


(B)

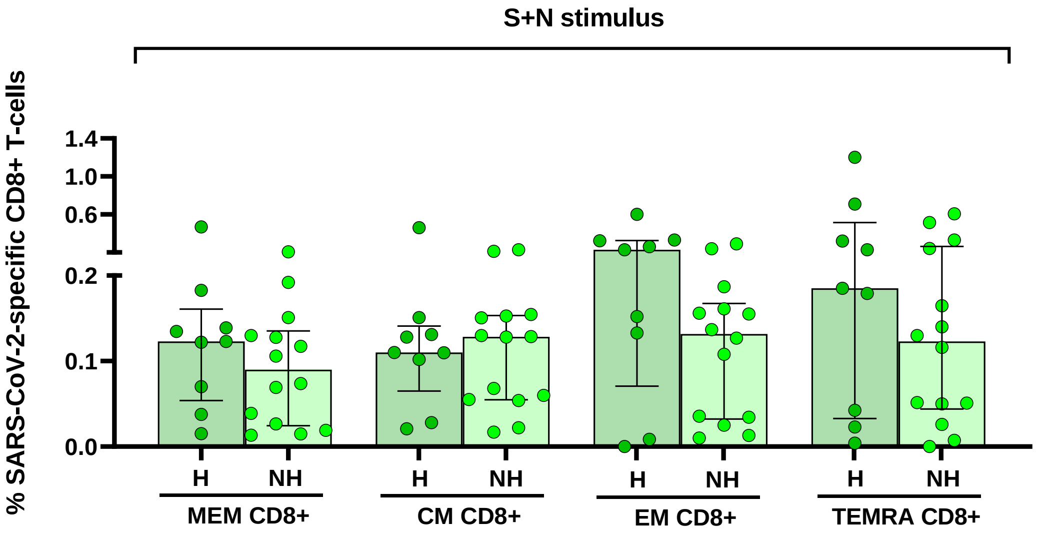


**Supplementary Figure 7. Combined SARS-CoV-2 specific T-cell response to N and S proteins in previously hospitalized and non-hospitalized subjects seven months after SARS-CoV-2 infection. (A)** Bar graphs represent the percentage of S plus N-specific CD4+ T-cell response. **(B)** Bar graphs represent the percentage of S plus N-specific CD8+ T-cell response. Each dot represents an individual. *p < 0.05, **p < 0.01, ***p < 0.001, ****p < 0.0001. Mann-Whitney U test was used for groups' comparisons.

**Supplementary Figure 8**

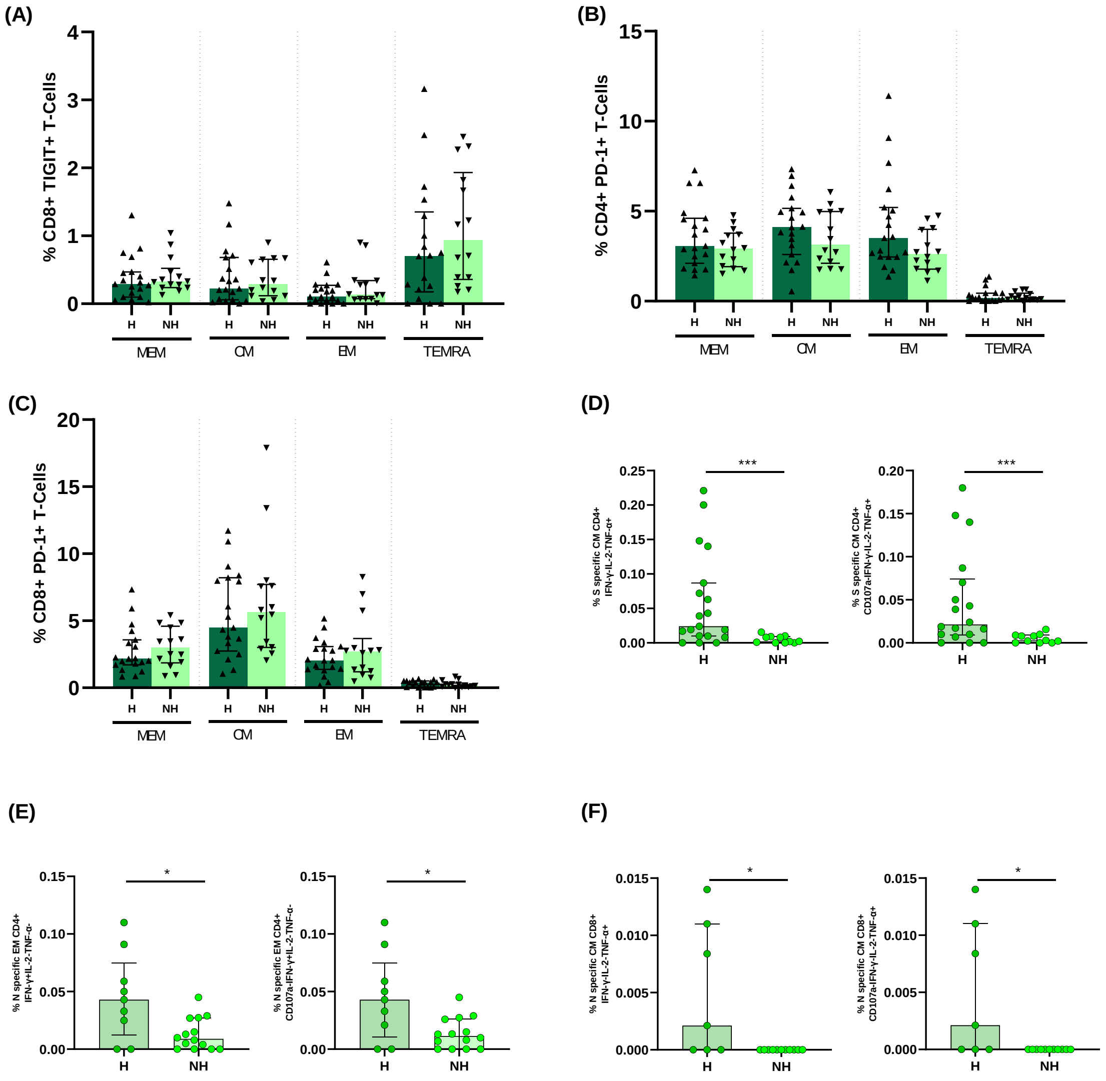


**Supplementary Figure 8. Additional features of T-cell exhaustion and SARS-CoV-2-specific T-cell response in previously hospitalized and non-hospitalized subjects seven months after SARS-CoV-2 infection. (A)** TIGIT expression in each CD8+ T-cell subset, **(B)** PD-1 expression in each CD4+ T-cell subset, **(C)** PD-1 expression in each CD8+ T-cell subset, in previously hospitalized and non-hospitalized subjects seven months after SARS-CoV-2 infection. **(D)** S-specific CM CD4+ T-cell levels of combinations only including TNF-α+ cells for three (IFN-γ, TNF-α, and IL-2) (left panel) and four (IFN-γ, TNF-α, IL-2 and CD107a) functions (right panel). **(E)** N-specific EM CD4+ T-cell levels of combinations only including INF-γ+ cells for three (IFN-γ, TNF-α, and IL-2) (left panel) and four (IFN-γ, TNF-α, IL-2 and CD107a) functions (right panel). **(F)** N-specific CM CD8+ T-cell levels of combinations only including TNF-α+ cells for three (IFN-γ, TNF-α, and IL-2) (left panel) and four (IFN-γ, TNF-α, IL-2 and CD107a) functions (right panel). The medians with the interquartile ranges are shown. Each dot represents an individual. $^{*}p < 0.05$, $^{**}p < 0.01$, $^{***}p < 0.001$, $^{****}p < 0.0001$. Mann-Whitney U test was used for groups' comparisons.

**Supplementary Figure 9**

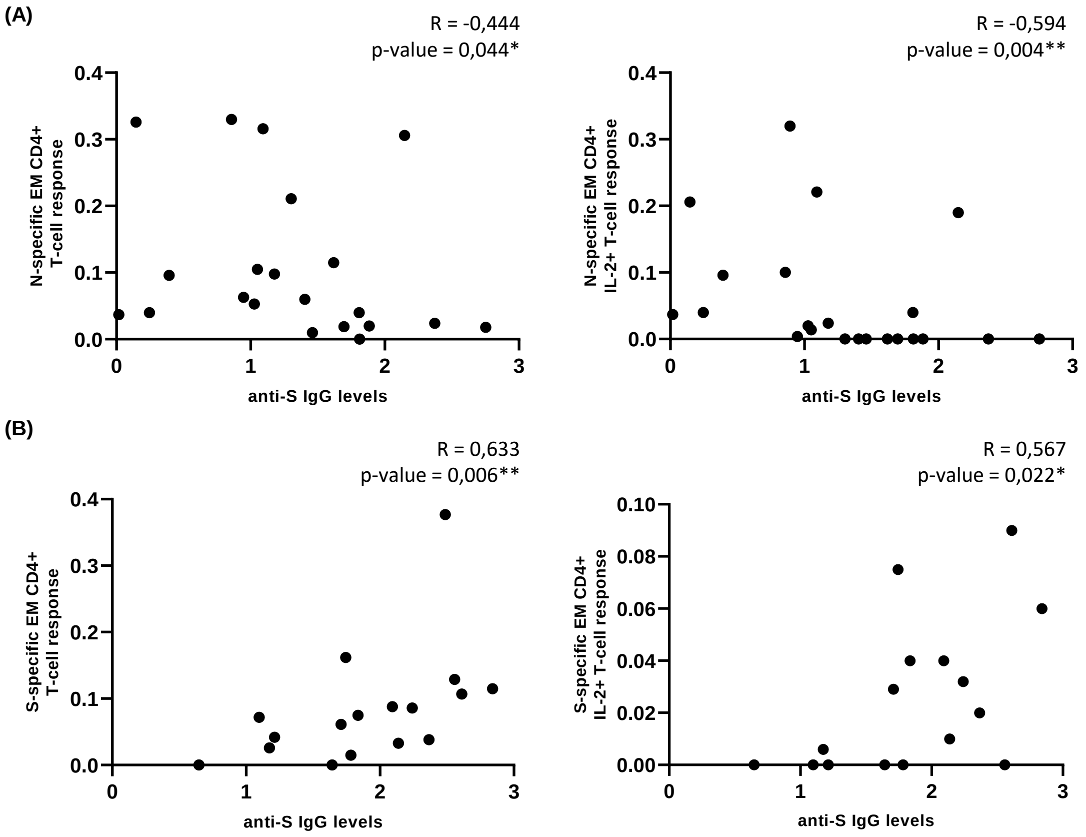


**Supplementary Figure 9. Anti-S IgG levels are directly correlated with S-specific T-cell response in acute infection but this correlation is inverse seven months after SARS-CoV-2 infection in previously hospitalized patients. (A)** Correlation graphs between anti-S IgG levels and the percentage of N-specific EM CD4+ T-cells (left panel) and N-specific CD4+ EM IL-2+ T-cells in acute SARS-CoV-2 infection. **(B)** Direct correlation between anti-S IgG levels and the percentage of S-specific EM CD4+ T-cells (left panel) and S-specific CD4+ EM IL-2+ T-cells in previously hospitalized patients seven months after SARS-CoV-2 infection (right panel). Each dot represents an individual. *p < 0.05, **p < 0.01, ***p < 0.001, ****p < 0.0001. Spearman test was used for non-parametric correlations.

**Supplementary Figure 10**

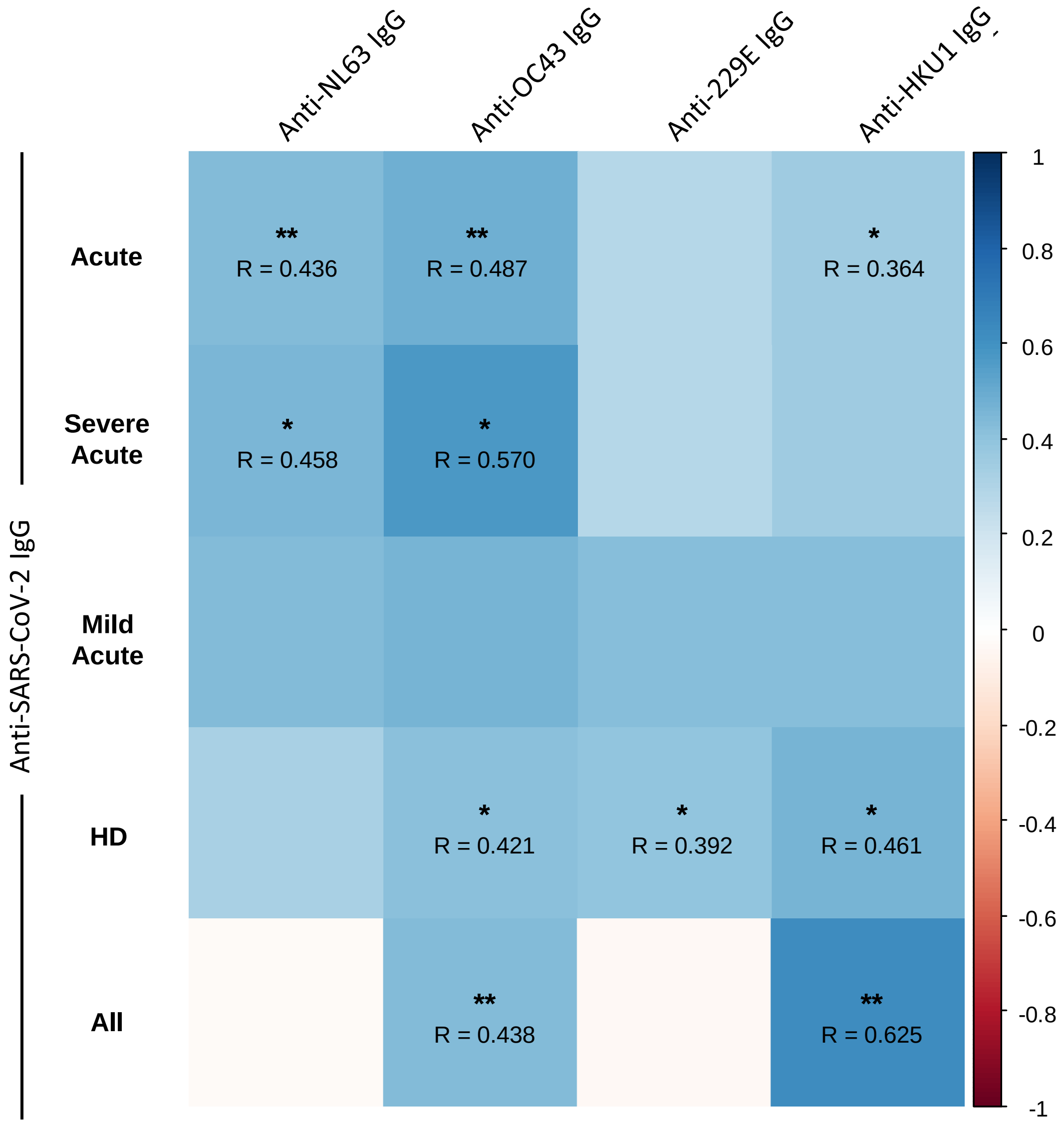


**Supplementary Figure 10. Anti-S IgG levels against endemic coronaviruses are associated with anti-S IgG levels against SARS-CoV-2.** Correlation matrix between anti-S IgG levels against SARS-CoV-2 and endemic coronaviruses (NL63, OC43, 229E and HKU1). Blue color represents positive correlations and red color shows negative correlations. The intensity of the color indicates the R coefficient. *$p < 0.05$, **$p < 0.01$, ***$p < 0.001$. Spearman test was used for non-parametric correlations.

**Supplementary Figure 11**

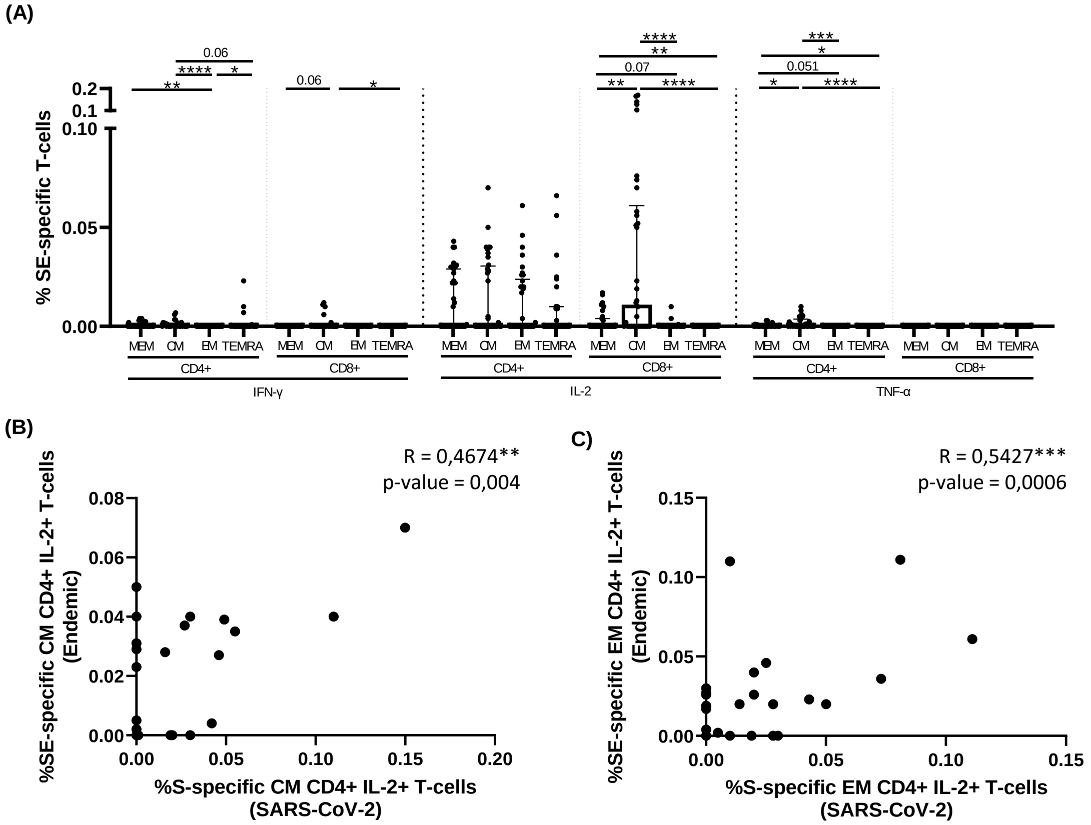


**Supplementary Figure 11. S-specific T-cell response in HD to endemic coronaviruses is mainly mediated by IL-2 production. (A)** Bar graph represent the SE-specific T-cell response in each T-cell subset for each cytokine. **(B)** Correlation between S-Specific and SE-specific CM CD4+ IL-2+ T cell levels and **(C)** Correlation between S-Specific and SE-specific EM CD4+ IL-2+ T cell levels. Each dot represents an individual. *p < 0.05, **p < 0.01, ***p < 0.001, ****p < 0.0001. Mann-Whitney U test was used for groups' comparisons and Spearman test for non-parametric correlations.

**Supplementary Figure 12**

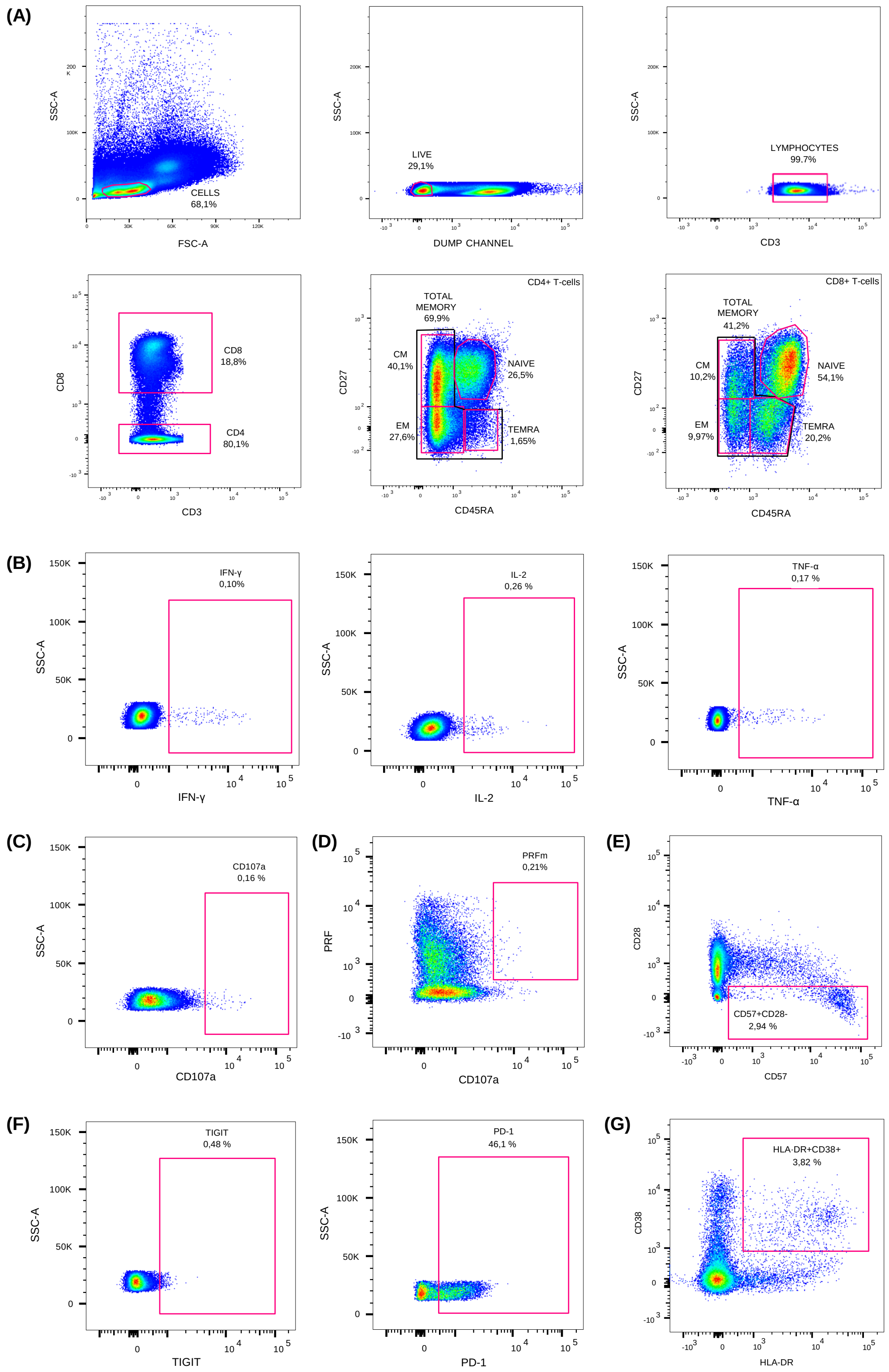


**Supplementary Figure 12. Schematic diagram of the gating strategy. (A)** Phenotyping of CD4+ and CD8+ T-lymphocyte subsets, including naïve (NAIVE); Total Memory (MEM), Central Memory (CM), Effector Memory (EM) and terminally differentiated memory (TEMRA) T cells. **(B)** Gating of intracellular cytokine production, including interferon gamma (IFN-γ), interleukin-2 (IL-2) and tumor necrosis factor alpha (TNF-α). **(C)** Degranulation factor, CD107a. **(D)** Cytolytic enzime, mature perforin (PRF). **(E)** Senescence, CD57+CD28-. **(F)** Exhaustion markers, T cell immunoreceptor with Ig and ITIM domains (TIGIT) and programmed death 1 molecules (PD-1) and **(G)** Activation, HLA-DR+CD38+.

### Supplementary Table 1. Characteristics of the study subjects.

| | Acute Infection | | | Discharged (6-8 months after diagnosis) | | | Healthy Donors | | |
|---|---|---|---|---|---|---|---|---|---|
| | All (n=37) | Mild (n=18) | Severe (n=19) | All (n=33) | Previously Hospitalized (n=19) | Previously Non Hospitalized (n=14) | All (n=33) | Old HD (n=19) | Young HD (n=14) |
| Age (years) | 71 [61.5 – 90] | 70 [57.75 – 76.5] | 72 [63 – 77] | 66 [58 – 74] | 71 [59 – 77] | 65.5 [56.75 – 71.5] | 62 [38.5 – 87] | 84 [71 – 90] | 38 [33.75 – 40.75] |
| Sex (Female sex), n (%) | 13 (35.1) | 6 (33.3) | 7 (36.8) | 12 (36.4) | 5 (26.3) | 7 (50) | 14 (42.4) | 10 (52.6) | 4 (28.6) |
| Oxygen Saturation ($SatO_2$), (%) | 93 [91 – 97] | 96 [93 – 98.25] | 91 [87 – 95] | N/A | N/A | N/A | N/A | N/A | N/A |
| Time since hospitalization, (days) | 3 [2 – 21.5] | 2.5 [1 – 3.25] | 18 [3 – 28] | 201 [180.5 – 221] | 187 [173 – 193] | 221 [218 – 230.75] | N/A | N/A | N/A |
| Time since symptoms onset, (days) | 17 [7 – 31.5] | 7.5 [4 – 19.5] | 31 [17 – 37] | 208 [190 – 232] | 186 [195 – 202] | 232 [224.5 – 239.25] | N/A | N/A | N/A |
| Time hospitalized, (days) | 16 [7.5 – 29] | 8 [6.5 – 12.25] | 26 [17 – 36] | N/A | 19 [9 – 37] | 0 | N/A | N/A | N/A |
| Comorbidities, n (%) | 29 (78.4) | 15 (83.3) | 14 (73.7) | 19 (57.6) | 15 (78.9) | 3 (21.4) | N/A | N/A | N/A |
| Diabetes mellitus | 12 (32.4) | 7 (38.9) | 5 (26.3) | 4 (12.1) | 3 (15.8) | 1 (7.1) | N/A | N/A | N/A |
| Hypertension | 25 (67.6) | 12 (66.7) | 13 (68.4) | 12 (36.4) | 10 (52.6) | 2 (14.3) | N/A | N/A | N/A |
| Cardiovascular disease | 11 (29.7) | 7 (38.9) | 4 (21.1) | 6 (18.2) | 4 (21.1) | 2 (14.3) | N/A | N/A | N/A |
| Obstructive pulmonary disease | 4 (10.8) | 2 (11.1) | 2 (10.5) | 0 | 0 | 0 | N/A | N/A | N/A |
| Malignancy | 4 (10.8) | 2 (11.1) | 2 (10.5) | 0 | 0 | 0 | N/A | N/A | N/A |
| Symptoms at admission (%) | | | | | | | | | |
| Cough | 24 (64.9) | 11 (61.1) | 13 (68.4) | 22 (66.7) | 12 (63.2) | 10 (71.4) | N/A | N/A | N/A |
| Fever | 27 (73) | 11 (61.1) | 16 (84.2) | 25 (75.8) | 14 (73.7) | 11 (78.6) | N/A | N/A | N/A |
| Dyspnea | 19 (51.4) | 7 (38.9) | 12 (63.2) | 16 (48.5) | 10 (52.6) | 6 (42.9) | N/A | N/A | N/A |
| Anosmia | 5 (13.5) | 0 | 5 (26.3) | 6 (18.2) | 6 (31.6) | 0 | N/A | N/A | N/A |
| Diarrhoea | 6 (16.2) | 3 (16.7) | 3 (15.8) | 9 (27.3) | 7 (36.8) | 2 (14.3) | N/A | N/A | N/A |
| Muscle pain | 3 (8.1) | 1 (5.6) | 2 (10.5) | 6 (18.2) | 5 (26.3) | 1 (7.1) | N/A | N/A | N/A |
| Treatment during hospitalization; n (%) | | | | | | | | | |
| Hydroxychloroquine | 33 (89.2) | 15 (83.3) | 18 (94.7) | N/A | 19 (100) | N/A | N/A | N/A | N/A |
| Lopinavir/Ritonavir | 24 (64.9) | 8 (44.4) | 16 (84.2) | N/A | 16 (84.2) | N/A | N/A | N/A | N/A |
| Beta Interferon | 12 (20) | 2 (11.1) | 10 (52.6) | N/A | 7 (36.8) | N/A | N/A | N/A | N/A |
| Corticoids | 18 (48.6) | 4 (22.2) | 14 (73.7) | N/A | 12 (63.2) | N/A | N/A | N/A | N/A |
| Remdesivir | 3 (8.1) | 3 (16.7) | 0 | N/A | 0 | N/A | N/A | N/A | N/A |
| Tocilizumab | 10 (27) | 1 (5.6Y) | 9 (47.4) | N/A | 10 (52.6) | N/A | N/A | N/A | N/A |

Categorical variables are expressed as number and percentages (%), and continuous variables are expressed as median (interquartile ranges [IQR]). N/A, not applicable. The different groups (acute infection, discharged patients and healthy donors) were age and sex matched. Chi-square test and a Mann-Whitney U test were used to compare categorical and continuous variables, respectively. Analysis by age; acute infection vs HD (p=0.423); discharged patients vs HD (p=0.700); Previously Hospitalized patients vs HD (p=0.635); Previously Non Hospitalized patients vs HD (p=0.907). Analysis by sex; acute infection vs HD (p=0.532); discharged patients vs HD (p=0.614); Previously Hospitalized patients vs HD (p=0.245); Previously Non Hospitalized patients vs HD (p=0.633). Severe participants were those who required Intensive Care Unit admission, or having ≥6 points in the ordinal scale score based on Beigel et al. (1) or death.
